# Non-Planar Perylene Diimides Dual-Targeting Mitochondrial Membrane Potential and Mitochondrial DNA Transcription for Antitumor Therapy

**DOI:** 10.1101/2025.10.03.680392

**Authors:** Junfeng Huo, Xin Jin, Meng Yin, Na Hu, Tianyu Liu, Wenwen Lu, Qingyi Tong, Jirong Huang, Huijuan You

**Affiliations:** Hubei Key Laboratory of Natural Medicinal Chemistry and Resource Evaluation, School of Pharmacy, Tongji Medical College, Huazhong University of Science and Technology, Wuhan 430030, China; College of Life and Environmental Sciences, Hangzhou Normal University, Hangzhou 311121, China; Key Laboratory of Organ Development and Regeneration of Zhejiang Province, Hangzhou 311121, China

**Keywords:** mitochondria, mitochondrial DNA transcription, OXPHOS inhibition, Breast Cancer, DNA intercalator

## Abstract

Mitochondrial DNA (mtDNA) transcription is an emerging target in cancer therapy. While enzyme-targeted inhibitors such as IMT1B block POLRMT, their organelle specificity remains unclear. Here, we report a distinct strategy using non-planar perylene diimide derivatives (PDIs). The fluorescent lead compound PDIC-BL combines mitochondrial membrane potential (ΔΨm) dependent accumulation (validated in cells and isolated mitochondria), and with direct intercalation into mtDNA (*K*_d_ = 32 μM), as demonstrated by single-molecule stretching assays. This dual-targeting mechanism leads to selective inhibition of mitochondrial transcription, ROS generation and triggers apoptosis. PDIC-BL exhibits superior antitumor efficacy in breast cancer cell lines (MCF-7, MDA-MB-231) and significantly supresses tumor growth in vivo, with good tolerability in cancer xenograft model in nude mice. This work not only elucidates the structure-activity relationship of non-planar PDIs, but also offers a generalizable strategy for developing organelle-selective DNA binding therapeutics.

## INTRODUCTION

Mitochondria are essential organelles responsible for key cellular functions, including oxidative phosphorylation (OXPHOS), metabolic regulation, and the control of apoptosis. Mitochondrial dysfunction has been implicated in various human diseases, including cancer, cardiovascular disorders, and neurodegeneration. Increasing evidence suggests that mitochondria play a pivotal role in malignant transformation and that tumor cells exhibit metabolic dependencies compared to their normal counterparts ^1^.

Human mitochondrial DNA (mtDNA) is a small, circular double-stranded DNA molecule of approximately 17,000 base pairs, encoding essential genes for OXPHOS machinery ^2-3^. Due to the endosymbiotic origin of mitochondria, most mitochondrial genes were relocated to the nuclear genome ^4^. Only 37 essential genes remain in the mitochondrial genome, encoding 13 proteins, 2 ribosomal RNAs, and 22 transfer RNAs, required for mitochondrial function and energy production ^2^. The transcription of these mitochondrially encoded genes is paramount for maintaining mitochondrial homeostasis, rendering mitochondria particularly sensitive to transcription inhibitors.

Targeting mtDNA transcription has recently emerged as a promising therapeutic target in cancer ^5-10^. Most current approaches focus on inhibiting mitochondrial RNA polymerase (POLRMT), the central enzyme responsible for mtDNA transcription, as demonstrated by small molecule allosteric inhibitors such as IMT1B ^6, 8-9^. Alternative strategies, such as proteolysis-targeting chimeras (PROTACs), and activation of mitochondrial matrix caseinolytic protease (ClpP) have also been used to degrade POLRMT for antitumor effects ^11-12^. Notably, contrary to the classical Warburg hypothesis, OXPHOS remains active or even upregulated in certain tumors, including a subset of breast cancers ^13^. In such cases, disrupting OXPHOS and hindering energy supply in OXPHOS-dependent cancer cells ultimately can lead to cell death ^14^.

Despite its promising profile, IMT1B presents unresolved challenges. A key knowledge gap remains regarding its sub-cellular localization-specifically, whether it effectively accumulates in mitochondria. Additionally, IMT1B demonstrated insufficient antitumor activity in certain cancer cell lines, particularly in breast cancer models such as MCF-7 and MDA-MB-231, where IC_50_ value exceeds 30 μM ^10^. POLRMT mutations (e.g., L796Q, L816Q) can impair IMT1B drug binding and result in drug resistance ^15^. These limitations highlight the need of alternative strategies to effectively target mtDNA transcription in cancer.

In addition to directly targeting RNA polymerase by IMT1B, an alternative strategy to disrupt transcription involves the use of DNA-intercalating agents like actinomycin D^16^. These intercalators insert aromatic rings between adjacent base pairs and interfere with the RNA polymerase activity. However, most previously reported DNA-interacting ligands, such as doxycycline ^17^ and ciprofloxacin ^18^ primarily act on nuclear DNA and show limited mitochondria accumulation. Some efforts have been made to redirect DNA binding drugs to mitochondria, such as through conjugation of cisplatin conjugation with mitochondrial targeting moieties like triphenylphosphonium (TPP) ^19^. Nonetheless, achieving both mitochondria-specific accumulation and mtDNA-binding affinity remains challenging.

A defining feature of many cancer cells is their elevated mitochondrial membrane potential (ΔΨm) relative to that of normal epithelial cells ^20^. This membrane potential represents the electrochemical gradient generated by proton pumping through the respiratory chain complexes across the inner mitochondrial membrane. The resulting negative charge within the mitochondria matrix drives the inward transport of cations and outward movement of anions ^21^. Consequently, many lipophilic cations, such as cyanine-based dyes (e.g., JC-1) and rhodamine derivatives (e.g., Rhodamine 123, MitoTracker series) can accumulate in mitochondria. Small molecules like Gboxin exploit this mechanism to selectively targets tumor mitochondria over normal cells, inhibits OXPHOS, and suppresses tumor proliferations ^22^. Other ΔΨmtargeting compounds including TPP derivatives have been demonstrated potential therapeutic effects through the generation of reactive oxygen species (ROS) ^23-24^. However, these mitochondrial targeting scaffolds generally lack intrinsic DNA-binding capability, and their ability to inhibit mtDNA transcription has not been thoroughly investigated.

Perylene diimides (PDIs) represent a class of polycyclic aromatic hydrocarbons (PAHs) with a rigidly planar, conjugated skeleton ^25-26^. This inherent planarity enables them to intercalate into the base pairs of double-stranded DNA (dsDNA) ^27-28^. Conventional PDIs tend to localize within the nucleus, where they intercalate into nuclear DNA ^27-28^. In contrast, recent studies have shown that non-planar PDIs, incorporating twisted skeleton (e.g., PDIC-NC) can accumulate in mitochondria, where they trigger apoptosis via ROS generation ^29-31^. These findings raise an intriguing hypothesis: non-planar PDIs may uniquely combine mitochondrial localization and the ability to bind mtDNA, potentially making them selective mtDNA transcription inhibitors. However, to date, no studies have directly investigated the capacity of non-planar PDIs to interact with mtDNA.

Herein, we report the design, synthesis, and functional characterization of a series of four non-planar PDIs (Figure 1). The lead compound, PDIC-BL, demonstrates mitochondria-specific accumulation, ΔΨm-dependent uptake, and direct mtDNA intercalation, resulting in mtDNA transcription inhibition, ROS elevation, and robust antitumor activity both in vitro and in vivo. Collectively, this study establishes non-planar PDIs as new class of selective mtDNA transcription inhibitors, offering a promising framework for the development of cancer therapeutics.

**Figure 1.**
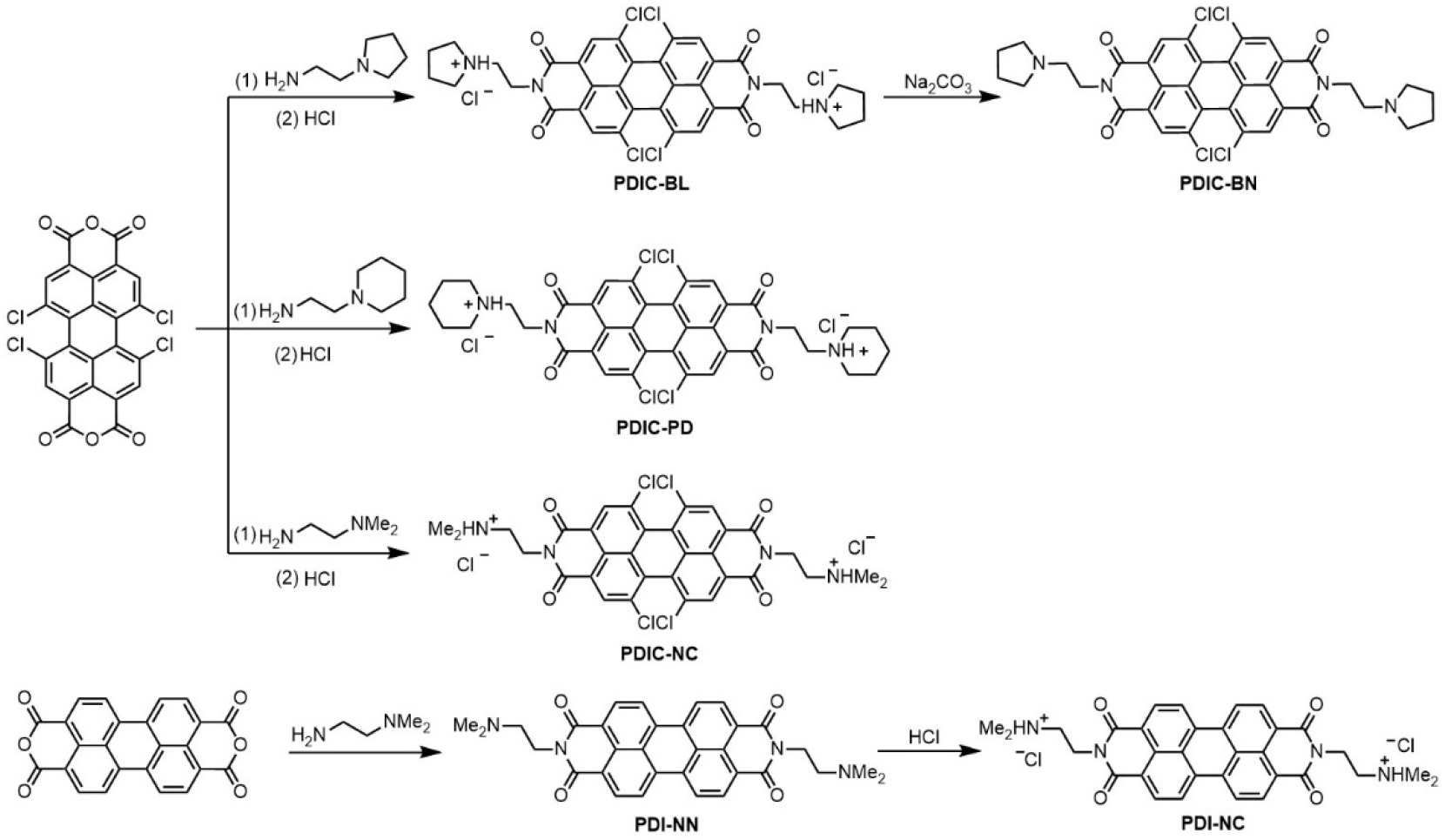
Synthesis of PDI derivatives. Synthesis route of non-planar perylene-1,6,7,12-tetrachloro-3,4,9,10tetracarboxyldiimide (PDIC) derivatives, including PDIC-BL, PDIC-BN, PDIC-PD and PDIC-NC. Planar PDI-NC was also synthesized for comparisons.

## RESULTS

### Non-planar Perylene Diimide Derivative PDIC-BL exhibit mitochondrial accumulation and potent antitumor activity

To explore how structural features influence the biological activity of PDIs, we designed and synthesized a series of non-planar PDIs: PDIC-BL, PDIC-BN, PDIC-PD, PDIC-NC, as well as the planar analogue PDI-NC (Figure 1). The non-planar compounds share a common 1,6,7,12-tetrachloro-substituted perylene core, where chlorination at the bay positions induces steric hindrance, disrupting π–π stacking and introducing a twisted, non-planar geometry. Detailed synthetic procedures and compound characterization by ^1^H NMR, ^13^C NMR, HRMS are provided in the supporting information (Figure S1-S17). This chlorination at the bay positions induces steric distortion of the perylene core significantly improved solubility in DMSO, which is critical for biological testing (Figure S18). Each PDIC derivative carries two side chains introduced via imide condensation with amine precursors, allowing for systematic variation in side-chain size and charge.

The antiproliferative activities of these compounds were then evaluated in MCF-7 and MDA-MB-231 breast cancer cells using the MTT assay (Figure 2A-B, Table S1), revealing a clear structure-activity relationship (SAR). Among the five compounds, PDIC-BL, emerged as the most potent with an IC_50_ of 0.59 ± 0.09 µM and 0.48 ± 0.02 µM in MCF-7 cells and MDA-MB-231, respectively. The non-planar PDIC-NC also demonstrated strong activity, (IC_50_ = 0.8 ± 0.2 µM and 0.39 ± 0.02 µM, in MCF-7 and MDA-MB-231), while its planar counterpart PDI-NC was approximately four-fold less active (ICLL = 3.0 ± 1.3 µM), highlighting the importance of non-planarity. The presence of a cationic side chain was essential for cytotoxicity: PDIC-BN, a neutral analogue of PDIC-BL, was nearly inactive (IC_50_ > 10 µM), underscoring the role of electrostatic interactions in cellular uptake and activity. Moreover, PDIC-PD, bearing a bulkier piperidinium group, was significantly less active (IC_50_ = 5.0 ± 1.0 µM), indicating that side-chain sterics fine-tune potency.

**Figure 2.**
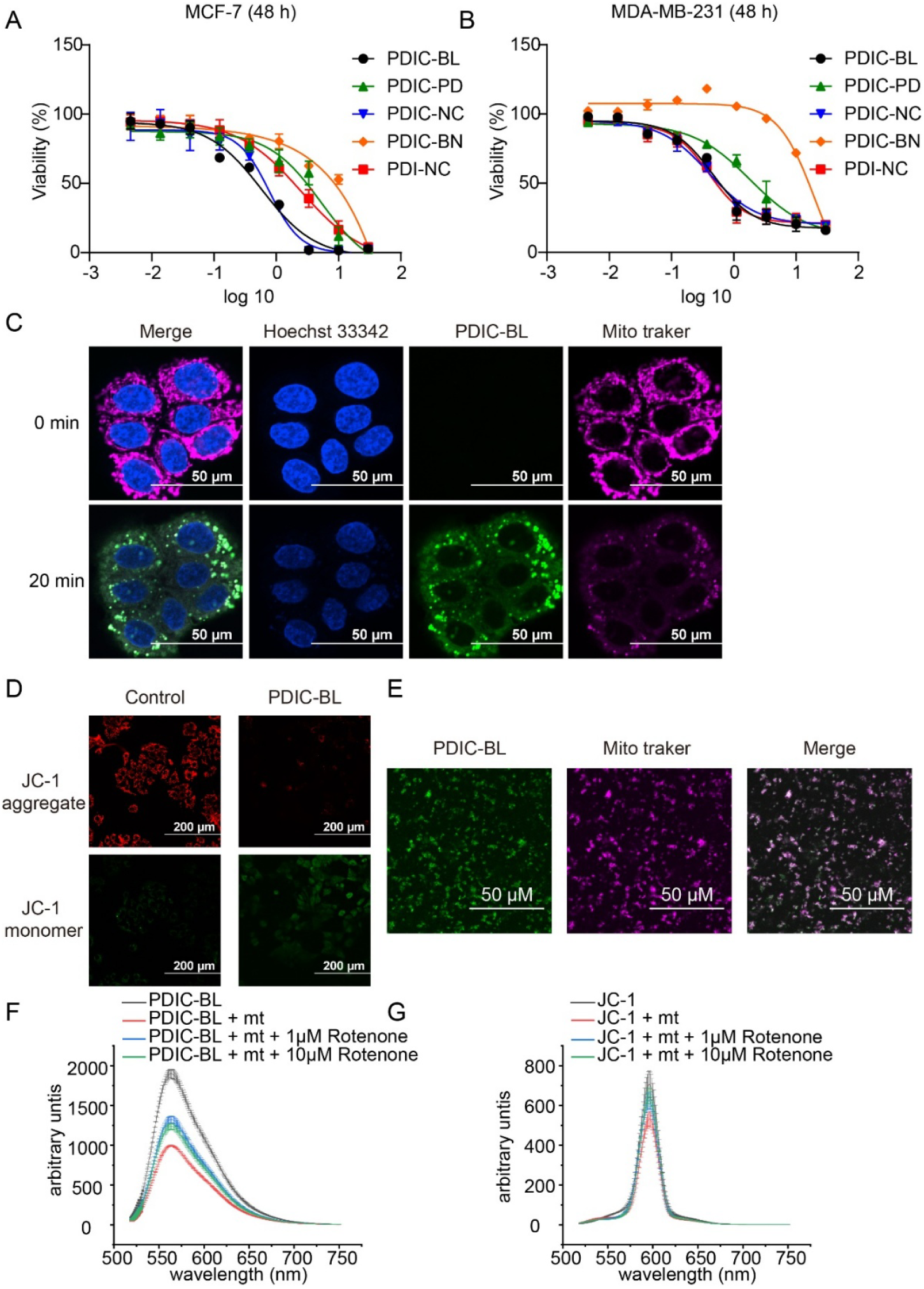
PDIC-BL demonstrates antitumor activity and mitochondrial accumulation. (A-B) PDIC derivatives inhibit the proliferation of breast tumor cell lines MCF-7 (A) and MDA-MB-231 (B). Dose-response cell viability curves were evaluated using the MTT assay after 48 hours of treatment. The calculated IC_50_ values were shown in Table S1. (C) Mitochondrial localization of PDIC-BL in MCF-7 cells. Confocal microscopy images show subcellular distribution of PDIC-BL (2 µM) in MCF-7 cells at 0 (before treatment) and 20 minutes (post-treatment). PDIC-BL rapidly accumulates in mitochondria with minimal nuclear localization. Cells were co-stained with Hoechst 33342 (nuclei, blue) and MitoTracker Deep Red FM (mitochondria, magenta). (D) PDIC-BL induces mitochondrial membrane depolarization. JC-1 assay demonstrates loss of mitochondrial membrane potential after 6 hours of PDIC-BL (2 µM) treatment, indicated by a shift from red fluorescent JC-1 aggregates to green fluorescent JC-1 monomers. (E) Live confocal imaging of isolated mitochondria confirms co-localization of PDIC-BL with mitochondrial structures. MitoTracker Deep Red FM (magenta) stains mitochondria; 2 µM PDIC-BL shows intrinsic green fluorescence. (F) Fluorescence quenching of PDIC-BL upon accumulation in isolated mitochondria. The fluorescence of PDIC-BL (10 µM, black curve) is quenched upon incubation with energized mitochondria (2.5 mg/mL, red curve). Pretreatment with rotenone (1 µM or 10 µM, blue and green curves, respectively) prevents this quenching by blocking mitochondrial uptake. (G) ΔΨm-dependent fluorescence quenching of the control dye JC-1. The fluorescence of JC-1 (1 µM, black curve) is similarly quenched by energized mitochondria (1.25 mg/mL, red curve), and this effect is blocked by pretreatment with rotenone (1 µM or 10 µM, blue and green curves).

Given its superior activity, PDIC-BL was selected for further investigations. It exhibited low IC_50_ value in cancer cell line HEK-293T, HCT-116 with a low IC_50_ value (<2 μM; Figure S19). Importantly, PDIC-BL showed lower toxicity toward primary mouse lymphocytes (3.0 ± 1.0 µM), suggesting partial selectivity for cancer cells (Figure S20).

We next investigated the subcellular localization of PDIC-BL. The compound exhibits strong intrinsic fluorescence with concentration-dependent emission peak at 565 nm (Figure S21), allowing for direct live-cell imaging. Confocal microscopy analysis, using the intrinsic fluorescence of PDIC-BL and the co-localization with mitochondria-specific dye MitoTracker Deep Red revealed that PDIC-BL rapidly accumulates in the mitochondria of MCF-7 cells (Figure 2C, Figure S22), with minimal nuclear accumulation. Quantitative analysis yielding a Pearson’s correlation coefficient of 0.80 between the PDIC-BL and MitoTracker signals, indicating a high degree of colocalization (Figure S23A). This contrasts with planar PDIs (PDI-NC), which have been reported to predominantly accumulate within the cell nucleus ^27^.

This mitochondrial accumulation coincided with a loss of mitochondrial membrane potential (ΔΨm), as evidenced by an increase in green JC-1 monomers after PDIC-BL treatment (2 µM, 6 hours), indicating mitochondrial membrane depolarization (Figure 2D).To directly assess whether ΔΨm drives the accumulation of PDIC-BL, we performed in vitro uptake assays using isolated mitochondria (Supporting information)^32^. The colocalization PDIC-BL and MitoTracker was first confirmed by confocal fluorescence imaging, with a Pearson’s coefficient of 0.91 (Figure 2E, Figure S23B). Moreover, upon the addition of energized mitochondria to a solution of PDIC-BL, its intrinsic fluorescence (λem = 565 nm) was significantly quenched, decreasing from 1960 ± 60 a.u. to 993 ± 7 a.u. (n =3)(Figure 2F). This quenching phenomenon is consistent with previously reported behaviors of the mitochondria-targeting dyes (JC-1, or rhodamine 123), which often undergo fluorescence decrease due to the formation of dimer or multimers within the mitochondrial matrix ^32^. Finally, isolated mitochondria were pre-depolarized with 1 µM rotenone (a complex I inhibitor) to induce membrane depolarization. Under these conditions, PDIC-BL fluorescence quenching was markedly attenuated, with emission intensity partially recovering to 1320 ± 40 a.u., indicating reduced mitochondrial accumulation in the absence of ΔΨm. A similar ΔΨm-dependent quenching effect was observed in a positive control experiment using the known cationic dye JC-1 (1 µM, λem = 595 nm) (Figure 2G).

Together, these results demonstrate that PDIC-BL exerts potent antitumor activity, driven by its non-planar, cationic structure, and that it selectively accumulates in mitochondria in a ΔΨm-dependent manner. This localization likely underpins its ability to induce mitochondrial dysfunction, supporting its potential as a mitochondria-targeted chemotherapeutic agent.

### PDIC-BL Directly Interacts with Mitochondrial DNA and Inhibits Its Transcription

Previous studies have shown that perylene bisimides like PDI-NC can intercalate into nuclear dsDNA, thereby inhibiting cancer cell growth^27-28^. However, PDIC-BL possesses a twisted non-planar perylene core, which may weaken π-π stacking interaction with DNA base pairs. Therefore, our next objective was to determine whether PDIC-BL directly interacts with mtDNA.

To investigate the interactions between PDIC-BL and mtDNA, we employed a single-molecule stretching assay using a dsDNA segment derived from human mtDNA ^33^. Binding of small intercalators to dsDNA leads to an increase in dsDNA contour length, providing a sensitive readout for detecting even weak DNA intercalators ^33-35^. The experimental design using magnetic tweezers was shown in Figure 3A, where a 5901 bp dsDNA segment was tethered between a paramagnetic bead and a cover glass slide, serving as a sensor to detect the DNA interacting agents. Figure 3B shows the force-extension curves of dsDNA measured in the presence of different concentrations of PDIC-BL at forces ranging from 1 pN to 75 pN. PDIC-BL induced a clear, concentration- and force-dependent elongation of dsDNA, consistent with classical intercalative binding behavior. In contrast, the non-cationic analogue, PDIC-BN showed negligible elongation under the same conditions (Figure 3C). This difference correlates with the weaker cellular activity of PDIC-BN, highlighting the importance of PDIC-BL’s cationic charge for DNA binding. Furthermore, a comparison with the planar derivative PDI-NC revealed that it produced even greater DNA elongation than the non-planar PDIC compounds (Figure S24), indicating that bay-region chlorination–induced twisting of the perylene core reduces intercalative capacity.

**Figure 3.**
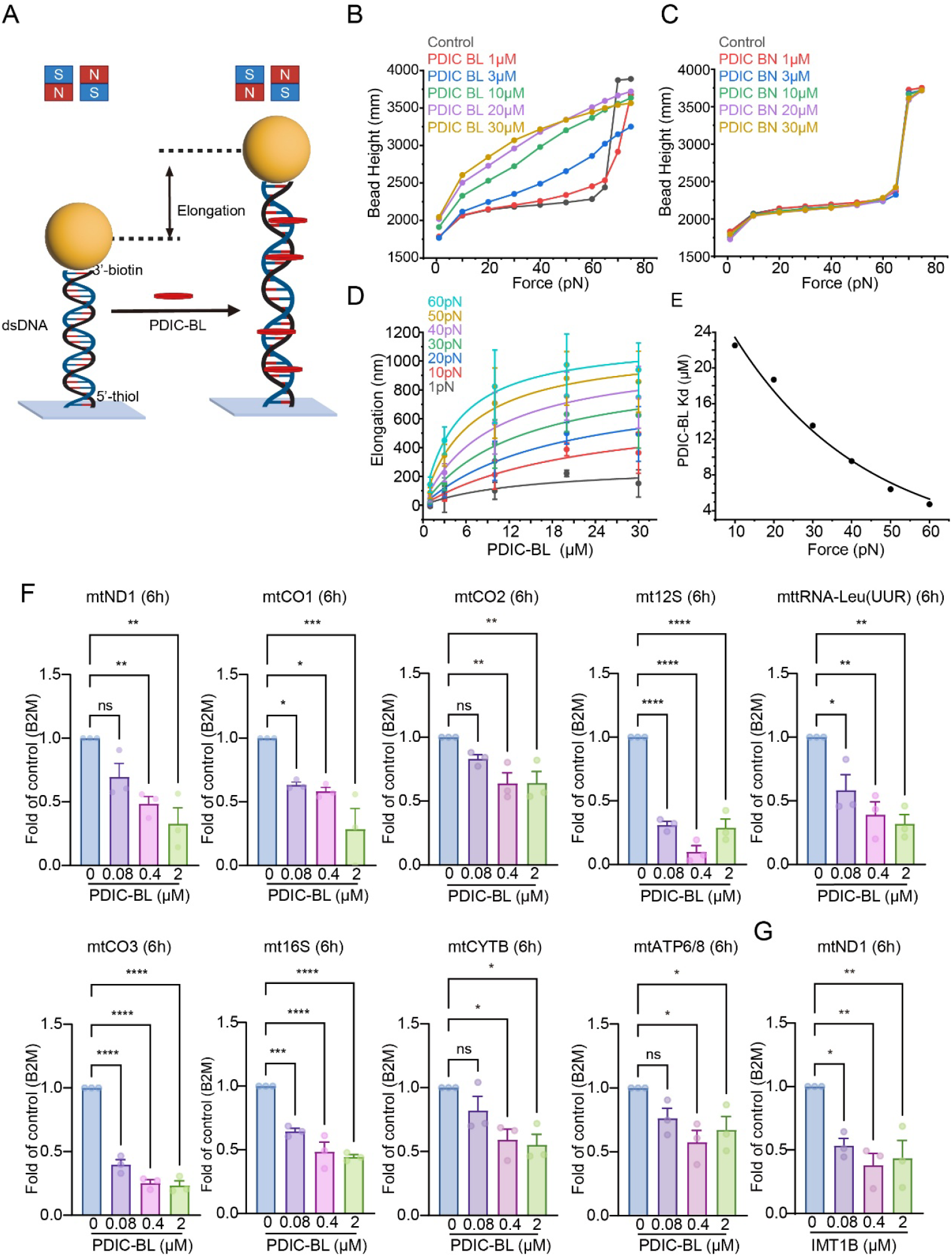
PDIC-BL intercalates into mtDNA and inhibits its transcription. (A) Schematic of the single-molecule stretching assay. A 5901 bp dsDNA with 5’-biotin and 5’-thiol label was tethered between a paramagnetic bead and a coverslip. (B) Force-extension curves for a single dsDNA molecule in the presence of varying concentrations of PDIC-BL (1 µM to 30 µM). The concentration-dependent increase in DNA length is a characteristic signature of DNA intercalation. (C) Force-extension curves for dsDNA in the presence of the neutral analogue, PDIC-BN (1 µM to 30 µM). (D) Elongation of dsDNA as a function of PDIC-BL concentration at forces ranging from 1 pN to 60 pN. Data are presented as mean ± SD, indicating greater elongation at higher forces and concentrations. (E) Dissociation constant (Kd) of PDIC-BL binding to dsDNA plotted against applied force, showing force-enhanced binding affinity. (F) Quantitative RT-PCR analysis of mitochondrial gene expression (e.g., mtND1, mtCO1, mtCO2, mt12S, mttRNA-Leu, mtCO3, mt16S) in cells treated with PDIC-BL at different concentrations (0, 0.08, 0.4, 2 µM) for 6 hours, showing significant, dose-dependent inhibition of transcription. Statistical significance: *p < 0.05, **p < 0.01, ***p < 0.001, ****p < 0.0001; ns, not significant. (G) For comparison, qRT-PCR analysis of mtND1 expression in cells treated with the POLRMT inhibitor IMT1B (0, 0.08, 0.4, 2 µM) for 6 h. Statistical significance: *p < 0.05, **p < 0.01.

To quantify the DNA binding affinity of PDIC-BL, the dissociation constants *K*_d_(*F*) at each applied force were calculated by fitting the data to the Hill equation ^36-37^, 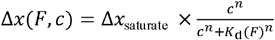, where Δ*x*(*F, c*) is the elongation of dsDNA at ligand concentration *c* and force *F*, Δ*x*_saturate_ is the saturating dsDNA elongation *n* is the Hill coefficient (Figure 3D). To estimate the *K*_d_ in the absence of force, the *K*_d_(*F*) values were fitted to the exponential force-dependent function, 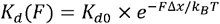, where k_B_ is the Boltzmann constant, and *T* is absolute temperature. The *K*_d_ for PDIC-BL was determined to be 32 ± 2 µM, which is comparable to that of several known DNA binding chemotherapeutic agents ^33^.

We next evaluated whether this binding correlates with mtDNA transcriptional inhibition using qRT-PCR. In MCF-7 cells, PDIC-BL induced broad and dose-dependent suppression of mitochondrial gene transcription, including transcripts for subunits of all respiratory complexes (e.g., mtND1, mtCO1, mtCO2), mitochondrial ribosomal RNAs (mt12S, mt16S) and a tRNA (Leu (UUR)) (Figure 3F). The most pronounced repression was observed for the 12S and 16S ribosomal RNA. This broad suppression of mtDNA-encoded gene expression corroborates the single-molecule data, indicating that PDIC-BL’s binding for mtDNA directly interferes with the mitochondrial transcription machinery.

To further establish a structure-activity relationship for mtDNA transcription inhibition, we also evaluated the effects of other key analogues. PDIC-NC, which also accumulates in mitochondria and exhibit strong cytotoxicity, significantly suppressed mtDNA transcription (Figure S25). In contrast, the bulkier and less potent PDIC-PD showed much weaker inhibitory effects (Figure S26) and planar analogue PDI-NC, which poorly localizes to mitochondria, showed little or no mtDNA transcription inhibition (Figure S27).

To benchmark PDIC-BL against known mitochondrial transcriptional inhibitors, we compared it with IMT1B, a selective POLRMT inhibitor. While both compounds broadly suppress mtDNA-encoded transcripts, PDIC-BL demonstrated stronger inhibition of mtCO1, mtCO3, ribosomal RNAs (mt12S rRNA, mt16S rRNA), and mttRNA-Leu UUR, but was less effective on mtATP6/8 and mtCYTB compared to IMT1B (Figure 3F, Figure S28). This distinct transcriptional profile suggests that PDIC-BL, as an mtDNA intercalator, inhibits transcription via a mechanism different from enzyme-targeting agents.

Given that the inhibition of mtDNA transcription by PDIC-BL can impair OXPHOS function, we next sought to determine if its cytotoxic effect is dependent on cellular metabolic states. To test this, we compared its antiproliferative activity in low-versus high-glucose media, as low-glucose conditions force cells to increase their reliance on OXPHOS. Indeed, the potency of PDIC-BL was significantly enhanced in MCF-7 and MDA-MB-231 cells under low-glucose conditions, with their respective IC_50_ values decreasing from 0.57 ± 0.06 µM to 0.11 ± 0.02 µM and from 1.15 ± 0.09 µM to ± 0.2 µM. In contrast, no significant change in cytotoxicity was observed in HCT-116 cells (Figure S29).

Together, these findings demonstrate that PDIC-BL directly binds to mitochondrial DNA, suppresses its transcription in a structure- and charge-dependent manner, and exerts enhanced cytotoxicity in cells reliant on mitochondrial metabolism.

### PDIC-BL Induces Reactive Oxygen Species Production and Apoptosis in the MCF-7 Cell Line

Having established that PDIC-BL inhibits mtDNA transcription, we next investigated the downstream cellular consequences. A hallmark of mitochondrial dysfunction is the over-production of reactive oxygen species (ROS). As hypothesized, treatment of MCF-7 cells with PDIC-BL (2 µM) led to a dramatic, dose-dependent increase in intracellular ROS levels, consistently observed through multiple detection methods (Figure 4A-B, Figure S30). Crucially, this effect was tightly linked to the key structural features responsible for its anticancer activity. Flow cytometry analysis revealed that the neutral analogue PDIC-BN (non-planar, neutral) and the planar analogue PDI-NC (planar, cationic) induced only marginal ROS production compared to the potent PDIC-BL (non-planar, cationic) (Figure 4C, 4D). This directly demonstrates that both the non-planar geometry, which enables mitochondrial accumulation, and the cationic side chains are essential for inducing severe oxidative stress.

**Figure 4.**
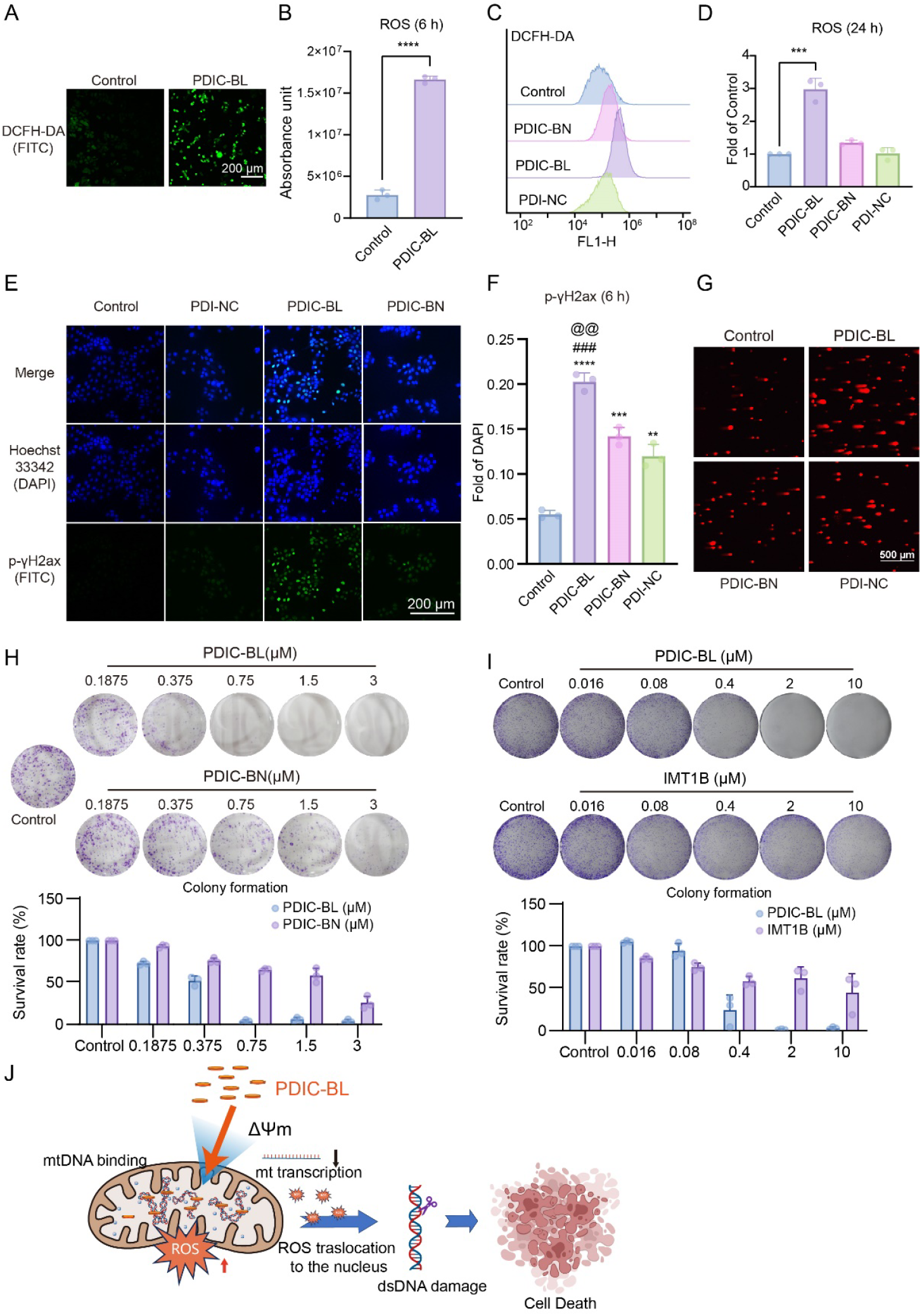
PDIC-BL induces mitochondrial ROS production, DNA damage and apoptosis in MCF-7 cell line. (A) Confocal microscopy showing reactive oxygen species (ROS) generation in MCF-7 cells treated with 2 µM PDIC-BL for 6 hours. ROS were visualized using DCFH-DA (green, FITC channel). (B) Quantification of fluorescence intensity from confocal images reveals a significant increase in intracellular ROS after PDIC-BL treatment (****p < 0.0001). (C) Flow cytometry analysis of ROS levels in MCF-7 cells treated with 1 µM PDIC-BL, PDIC-BN, or PDI-NC for 24 hours using DCFH-DA. (D) Statistical analysis of ROS level by flow cytometry after 24-hour treatment. PDIC-BL significantly increased ROS compared to PDIC-BN and PDI-NC (***p < 0.001). (E) Immunofluorescence images showing p-γH2AX foci formation, indicative of nuclear DNA damage, in MCF-7 cells treated with 2 µM PDIC-BL, PDIC-BN, or PDI-NC for 6 hours. (F) Quantification of γ-H2AX-positive nuclei. All treatment groups showed a significant increase in DNA damage compared to the control group. Notably, PDIC-BL induced the highest level of damage, which was significantly greater than that induced by both PDIC-BN (@@p < 0.01) and PDI-NC (###p < 0.001). Statistical significance is denoted as **p < 0.01, ***p < 0.001, and ****p < 0.0001. (G) Comet assay showing DNA strand breaks in MCF-7 cells treated with 1 µM PDIC-BL, PDIC-BN, or PDI-NC. PDIC-BL treatment induced the most significant comet tail formation. (H-I) Clonogenic survival assays in MCF-7 cells. (H) Cells were treated with PDIC-BL or PDIC-BN for 24 h, followed by recovery in drug-free medium for 7 days. (I) Cells were treated continuously with PDIC-BL or IMT1B for 7 days. Bar graphs quantify the survival rates relative to the untreated control. Data are presented as mean ± SD (n=3); statistical significance versus the control group is shown. (J) Proposed mechanism of action for PDIC-BL.

To investigate whether the observed ROS production was the primary cause of PDIC-BL-induced cytotoxicity, we pretreated MCF-7 cells with the potent antioxidant N-acetylcysteine (NAC) before exposure to the compound. Surprisingly, NAC pre-treatment failed to attenuate the substantial increase in intracellular ROS levels. Consistent with this, the cytotoxicity of PDIC-BL was not rescued by NAC, with no significant shift in the IC□□ value observed (Figure S31). This result suggests that while PDIC-BL potently induces ROS, this oxidative stress is likely a downstream consequence of severe and irreversible mitochondrial damage, rather than the primary, causative event leading to cell death.

To further explore the downstream consequences of mitochondrial dysfunction induced by PDIC-BL, we examined the integrity of nuclear DNA. Immunofluorescence staining for γ-H2AX, a well-established marker for dsDNA breaks, revealed a substantial increase in γ-H2AX foci in MCF-7 cells following PDIC-BL treatment (2 µM, 6 hours) (Figure 4E, 4F). In contrast, cells treated with structurally related PDIC-BN and PDI-NC showed only minimal γ-H2AX signals, which is consistent with their limited mitochondrial effects. The presence of dsDNA breaks was further confirmed by the comet assay, in which PDIC-BL induced the most prominent comet tails formation (Figure 4G). These observations suggest that the mitochondrial perturbations initiated by PDIC-BL— including membrane depolarization and transcriptional suppression—ultimately culminate in nuclear DNA damage.

To assess the long-term outcome of this damage cascade, we performed clonogenic survival assays. PDIC-BL significantly impaired the colony-forming ability of MCF-7 cells in a dose-dependent manner, reflecting irreversible loss of proliferative capacity. Crucially, its efficacy was markedly superior to both its structurally similar but inactive analogue PDIC-BN and the POLRMT inhibitor IMT1B (Figure 4H, Figure 4I). Collectively, these results demonstrate that PDIC-BL exerts potent and sustained antiproliferative effects through a mitochondria-centered mechanism involving transcriptional disruption, bioenergetic collapse, and nuclear DNA strand breaks, as illustrated in the proposed model (Figure 4J).

### PDIC-BL Demonstrates Potent Antitumor Efficacy with a Favorable Safety Profile in Vivo

Encouraged by the in vitro results, we assessed the therapeutic potential of PDIC-BL in an MCF-7 xenograft model using female BALB/c nude mice. Since IMT1B exhibited poor antiproliferative activity in breast cancer cell lines (IC_50_ > 30 µM, Figure S29D), we selected the clinically approved chemo-therapeutic agent doxorubicin (DOX) as a benchmark for antitumor efficacy. Mice were randomly assigned to three groups (n = 5 per group): control (saline), DOX (3 mg/kg/day), and PDIC-BL (2 mg/kg/day). All treatments were administered by daily intraperitoneal injections for 19 consecutive days. Both PDIC-BL and DOX treatment significantly inhibited tumor growth compared to the control group, with suppression becoming apparent from day 5 (Figure 5A). Notably, from day 11 onwards, PDIC-BL produced a significantly stronger antitumor effect than DOX (P < 0.05). After 19 days, average tumor volumes were 139 ± 77 mm^3^ for PDIC-BL, 242 ± 81 mm^3^ for DOX group and 769 ± 96 mm^3^ for the control group, confirming the superior efficacy of PDIC-BL (Figure 5B). These results were further supported by tumor weights (Figure 5C).

**Figure 5.**
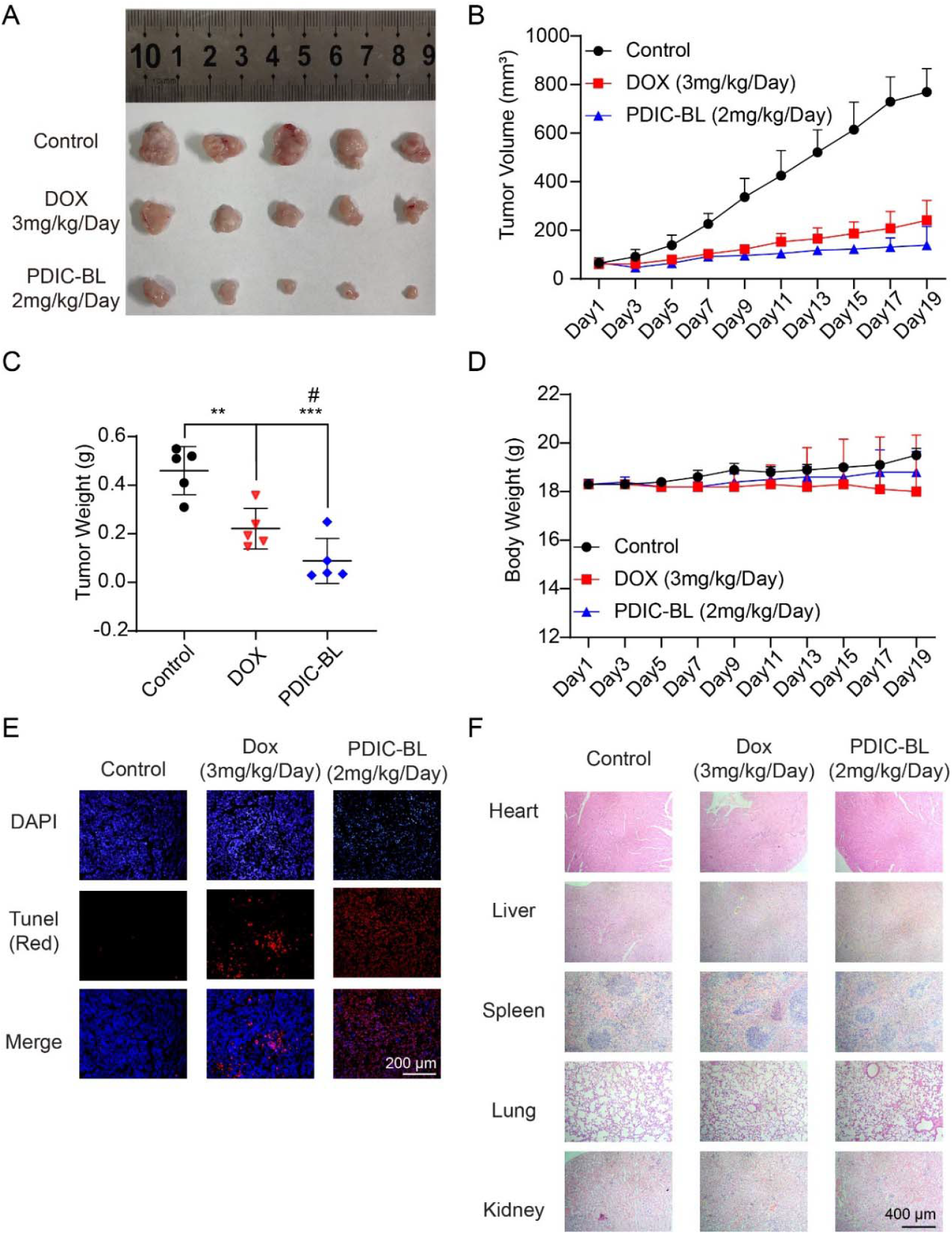
Therapeutic efficacy and safety of PDIC-BL in a breast cancer xenograft model. (A) Representative images of excised tumors from nude mice after 19 days of treatment with PDIC-BL (2 mg/kg/day), DOX (3 mg/kg/day), or vehicle control. (B) Tumor volume measurements over the 19-day treatment period. Data are presented as mean ± SD (n = 5 per group). PDIC-BL significantly suppressed tumor growth compared to the control and showed greater efficacy than DOX. (C) Tumor weight analysis at the end of the treatment period. PDIC-BL treatment resulted in significantly reduced tumor weight compared to both the control (***p < 0.001) and DOX groups (#p < 0.05). The DOX group also showed a significant reduction compared to the control (**p < 0.01). Final mean tumor weights (± SD) were: Control: 0.46 ± 0.10 g, DOX: 0.22 ± 0.08 g, and PDIC-BL:0.09 ± 0.09 g. (D) Body weight monitoring of mice during the treatment period, indicating no significant weight loss in any group, suggesting low systemic toxicity. (E) TUNEL staining of tumor sections showing increased apoptosis (red fluorescence) in the PDIC-BL-treated group compared to the control and DOX group. DAPI (blue) stains nuclei. (F) H&E staining of major organs (heart, liver, spleen, lung, and kidney) showed no significant histological abnormalities in the PDIC-BL group, indicating minimal off-target toxicity compared to DOX.

Importantly, PDIC-BL was well tolerated, with treated mice maintained stable body weight throughout the study, comparable to the control group. In contrast, mice treated with DOX exhibited a modest but statistically significant decrease in body weight loss (p = 0.073) suggesting mild systemic toxicity (Figure 5D).

To confirm that the observed antitumor effect was driven by the same mechanism identified in vitro, we analyzed the excised tumor tissues from the xenograft model. TUNEL staining revealed a significantly higher level of apoptosis in tumors from the PDIC-BL-treated group compared to both control and DOX group, confirming effective induction of cell death (Figure 5E). To assess mitochondrial transcriptional suppression, we performed qRT-PCR analysis on total RNA extracted from tumor tissues. While not all mitochondrial DNA encoded genes showed significant downregulation, key transcripts such as mtND1, mtCO2, and mt12S were suppressed by 52 ± 14%, 57 ± 11%, and 46 ± 14%, respectively, in PDIC-BL-treated tumors compared to controls, indicating a significant inhibition of mitochondrial transcription in vivo (Figure S32). Finally, to evaluate systemic toxicity, major organs including heart, liver, spleen, lung, and kidney were collected from each group for histological examination via H&E staining (Figure 5F). No histopathological abnormalities were observed in any treatment group, suggesting that PDIC-BL treatment does not induce overt tissue damage, and is well tolerated in vivo (Figure 5F). Overall, these results demonstrate that PDIC-BL retains its mitochondrial-targeting mechanism in vivo, effectively induces apoptosis, and exhibits favorable safety profile, highlighting its potential as a promising mitochondria-targeted anticancer agent.

## DISCUSSION

In this study, we developed a non-planar PDI derivative PDIC-BL to achieve dual-targeting of mitochondrial membrane potential ΔΨm and mtDNA transcription. Among the synthesized compounds, PDIC-BL emerged as the lead candidate, exhibiting stronger anti-proliferative activity in breast cancer cell lines (MCF-7 and MDA-MB-231) compared to the previously reported mitochondrial RNA polymerase inhibitor IMT1B ^6^. Our results clearly demonstrate that PDIC-BL selectively accumulates in mitochondria of breast cancer cell line MCF-7. Importantly, single-molecule stretching assay revealed that PDIC-BL can directly intercalate into mtDNA (*K*_d_ = 32 ± 2 μM), providing the direct biophysical evidence of DNA interaction by a non-planar PDI derivative. This intercalation leads to the inhibition of transcription of essential mitochondrial genes (tRNA, 12SRNA, 16SRNA), resulting in the disruption of mitochondrial function, increased mitochondrial reactive oxygen species (ROS) generation, and ultimately, the induction of apoptosis.

Interestingly, although both PDIC-BL and PDI-NC are cationic and structurally derived from perylene cores, their subcellular localizations differ significantly ^27-28^. This suggests that bay-region substitution may play roles in subcellular trafficking. Previous research hypothesized that the non-planar twisted perylene skeleton might hinder its DNA intercalation due to weakened π-π interactions^29^. However, our single-molecule magnetic tweezers assay revealed that PDIC-BL retains the ability to intercalate into dsDNA. Notably, although planar perylene PDI-NC showed stronger DNA intercalation (Figure S24), it showed negligible mtDNA transcription inhibition (Figure S27), highlighting that mitochondrial accumulation, not just intercalation strength, is essential for functional impact.

Beyond breast cancer, PDIC-BL also exhibited activity against other cancer cell lines such as HCT-116 and HEK-293T, highlighting the broader relevance of this approach across tumor types. Given that fast-growing tumor cells require mitochondrial metabolism, particularly OXPHOS to support biosynthesis and bioenergetic needs ^5^, targeting mtDNA transcription offers a selective anticancer strategy. Notably, normal cells such as mouse lymphocytes isolated from the spleen and lymph nodes showed less sensitivity to PDIC-BL (Figure S20). Moreover, PDIC-BL showed no significant toxicity in normal tissues (e.g., liver, heart) after 19 days of intraperitoneal injection of PDIC-BL (2 mg/kg) in mice, despite inducing a strong antitumor response in human cancer xenografts.

Mechanistically, PDIC-BL exerts its antitumor effects not only through mtDNA transcriptional inhibition but also by perturbing mitochondrial membrane potential and inducing oxidative stress (Figure 2D, Figure 4). This dual mode of action further differentiates PDIC-BL from IMT1B, which does not induce ROS production ^6^. JC-1 staining confirmed that PDIC-BL significantly perturbs ΔΨm, suggesting that its presence at the inner mitochondrial membrane may directly affect electron transport or proton flux. This aligns with previous research on perylene derivative (PDIC-NC)^29-31^, which were shown to generate ROS and reduce ATP production and oxygen consumption in A549 cells ^29^. These multifunctional disruptions of mitochondrial homeostasis likely contribute to the more potent anti-proliferative effects than IMT1B in MCF-7 and MDA-MB-231 cells (Figure S29). Moreover, mitochondrial ROS burst generated by sulfonated perylene PDIC-NS have been shown to trigger the mtDNA and nuclear DNA secretion into cytoplasm and activate the stimulator of interferon genes (STING) pathway, which is a promising immunotherapy strategy ^38^. Whether PDIC-BL triggers similar immune activation warrants further investigation.

Despite the good therapeutic activity, the genotoxic potential of DNA-binding compounds remains a critical consideration for clinical translation. The mitochondrial accumulation property of PDIC-BL may minimize exposure to nuclear DNA and thereby reduce carcinogenic risk ^39^. However, future studies are needed to assess genotoxicity of such compounds in both nuclear DNA and mtDNA, to evaluate their long-term safety, and to explore further structural refinements.

## ASSOCIATED CONTENT

## Supporting Information

Material and methods, including synthesis and characterization of perylene diimides derivatives, cell culture, isolation of crude mitochondrial fraction from cells, cytotoxicity assay, colony formation assay, comet assay, live-cell confocal microscopy, detection of intracellular reactive oxygen species, immunofluorescence staining for γ-H2AX, quantitative real-time PCR, nude mouse xenograft model, hematoxylin and eosin (H&E) staining, TUNEL staining, single molecule stretching assay, LC-MS analysis, ROS scavenging and cytotoxicity rescue assay, and additional Table S1-S3 and Figures S1−S32 as mentioned in the text. This material is available free of charge via the Internet at http://pubs.acs.org.

## AUTHOR INFORMATION

### Author Contributions

‡These authors contributed equally.

### Notes

The authors declare no competing financial interests.

## ACKNOWLEDGMENT

The authors thank Zhongwen Chen, Yifan Ge, Le Li, Jun Zhao for the stimulating discussions. The authors appreciate the help from technicians at the Analytical and Testing Centre of the Huazhong University of Science and Technology with the NMR measurements.

## ABBREVIATIONS

PDIs: Perylene diimides
OXPHOS: oxidative phosphorylation;

---

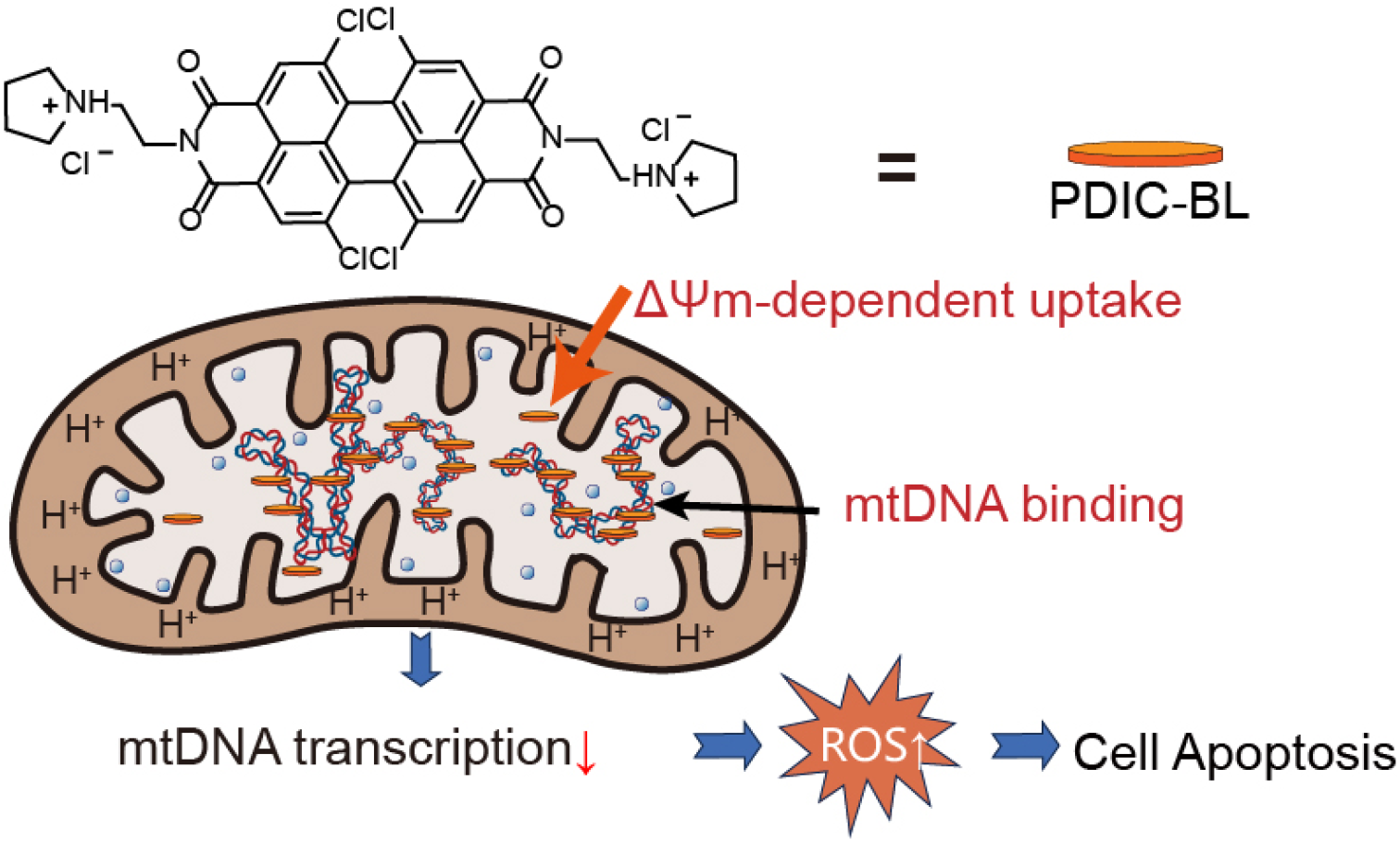

