## Supplemental Data 1 for "Non-Planar Perylene Diimides Dual-Targeting Mitochondrial Membrane Potential and Mitochondrial DNA Transcription for Antitumor Therapy"

Table of Content

Material and Methods

Supplementary Tables

**Table S1. IC_50_ values of compounds against MCF-7 and MDA-MB-231 breast cancer cells**

**Table S2. Primers used for mitochondrial gene single-molecule stretching assay**

**Table S3. Primers used for mitochondrial gene qPCR analysis**

Supplementary Figures

**Figure S1. The ^1^H NMR of PDIC-BL.**

**Figure S2. The ^13^C NMR of PDIC-BL.**

**Figure S3. The HRMS-ESI of PDIC-BL.**

**Figure S4. The ^1^H NMR of PDIC-BN.**

**Figure S5. The ^13^C NMR of PDIC-BN.**

**Figure S6. The HRMS-ESI of PDIC-BN**

**Figure S7. The ^1^H NMR of PDIC-PD.**

**Figure S8. The ^13^C NMR of PDIC-PD.**

**Figure S9. The HRMS-ESI of PDIC-PD.**

**Figure S10. The ^1^H NMR of PDIC-NC.**

**Figure S11. The ^13^C NMR of PDIC-NC.**

**Figure S12. The HRMS-ESI of PDIC-NC.**

**Figure S13. The ^1^H NMR of PDI-NC.**

**Figure S14. The ^13^C NMR of PDI-NC.**

**Figure S15. The HRMS-ESI of PDI-NC.**

**Figure S16. The ^1^H NMR of PDI-NN.**

**Figure S17. The ^13^C NMR of PDI-NN.**

**Figure S18. Solubility profiles of PDI derivatives under different solvent conditions.**

**Figure S19. Cytotoxicity of PDIC-BL in various cancer cell lines.**

**Figure S20. Cytotoxicity of PDIC-BL in mouse lymphocyte cells.**

**Figure S21. Concentration-dependent fluorescence emission of PDIC-BL**.

**Figure S22.** **Confocal imaging of PDIC-BL cellular uptake and comparison with doxorubicin.**

**Figure S23. Quantitative colocalization analysis of PDIC-BL and Mitochondria**.

**Figure S24.** **DNA binding properties of PDIs analyzed using single-molecule stretching assay.**

**Figure S25. PDIC-NC suppress mtDNA transcription in MCF-7 cells.**

**Figure S26.** **PDIC-PD suppress mtDNA transcription in MCF-7 cells.**

**Figure S27.** **PDI-NC exhibits negligible mtDNA transcription inhibition in MCF-7 cells.**

**Figure S28.** **IMT1B suppress mtDNA transcription in MCF-7 cells**.

**Figure S29. Enhanced cytotoxicity of PDIC-BL under low-glucose condition in cancer cells.**

**Figure S30. PDIC-BL induces dose-dependent ROS generation in MCF-7 Cells.**

**Figure S31**. **NAC pretreatment does not attenuate PDIC-BL induced ROS production or cytotoxicity.**

**Figure S32.PDIC-BL exhibits modest inhibition of mitochondrial gene transcription in vivo.**

**Material and Methods**

**1** **Synthesis and characterization of perylene diimides derivatives**

All nuclear magnetic resonance (NMR) data were recorded on a Bruker AM-600 spectrometer with TMS as the internal standard. The structures of PDIC-BL, PDIC-NC, PDI-NC were elucidated by analysis of their spectroscopic data (NMR and HRMS-ESI, Figures S1-S17).

*Synthesis of PDIC-BL and PDIC-BN:* 1,6,7,12-tetrachloro-3,4,9,10-tetracarboxylic dianhydride (265.0 mg 0.5 mmol) and 1-(2-aminoethyl) pyrrolidine (0.5 mL 3.9 mmol) were successively added into a mixed solvent of NMP (9 mL) and glacial acetic acid (8 mL) under argon protection. The black solid PDIC-BL (318.1 mg, 80%) precipitated by adding HCl (2 M, 20 mL) to the reaction after stirring at 120 °C for 24 h. ^1^H NMR (600 MHz, CF_3_COOD) δ 9.18 (s, 4H), 8.49 (s, 2H), 5.19 (t, *J* = 5.5 Hz, 4H), 4.53 (dd, *J* = 11.3, 5.9 Hz, 4H), 4.22 (q, *J* = 5.6 Hz, 4H), 3.71 (dt, *J* = 12.1, 6.3 Hz, 4H), 2.73-2.63 (m, 8H). ^13^C NMR (151 MHz, CF_3_COOD) δ 161.07, 132.64, 130.19, 127.17, 126.32, 119.15, 117.68, 52.06, 50.92, 33.27, 18.67. The compound PDIC-BL was dissolved in deionized water, and the resulting solution was treated with 20 mL of 5% aqueous sodium carbonate for 24 h. N,N’-Bis(2-(1-Pyrrolidinyl)ethyl)perylene-1,6,7,12-tetrachloro-3,4,9,10-tetracarboxyldiimide (PDIC-BN) (216.7 mg, 75%) was produced as a dark red solid. ^1^H NMR (600 MHz, CF_3_COOD) δ 9.17 (s, 4H), 5.18 (t, *J* = 5.7 Hz, 4H), 4.49 (dt, *J* = 11.9, 5.7 Hz, 4H), 4.20 (q, *J* = 5.7, 4.8 Hz, 4H), 3.75-3.69 (m, 4H), 2.72-2.60 (m, 8H). ^13^C NMR (151 MHz, CF_3_COOD) δ 161.04, 132.80, 130.35, 127.26, 126.44, 119.20, 117.63, 52.21, 51.19, 33.35, 18.78. HRMS-ESI, m/z calcd for C_36_H_29_Cl_4_N_4_O_4_^+^: 723.0908; found: 723.0904[M+H]^+^.

*Synthesis of PDIC-PD:* A mixture of 1,6,7,12-tetrachloro-3,4,9,10-tetracarboxylic dianhydride (265 mg, 0.5 mmol), 1-(2-aminoethyl) piperidine (0.5 mL, 3.5 mmol), 6 mL of glacial acetic acid, and 9 mL of NMP reacted under argon protection. After stirring at 120 °C for 24 h, N,N’-Bis(2-(1-piperidino)ethyl)perylene-1,6,7,12-tetrachloro-3,4,9,10-tetracarboxyldiimide hydrochlorate (PDIC-PD) (329.3 mg, 80%) precipitated as a dark red solid by adding HCl (2 M 10 mL) to the reaction. ^1^H NMR (600 MHz, CF_3_COOD) δ 9.20 (s, 4H), 7.75 (s, 2H), 5.21 (t, *J* = 5.6 Hz, 4H), 4.44-4.38 (m, 4H), 4.12 (q, *J* = 5.5 Hz, 4H), 3.50 (q, *J* = 11.4 Hz, 5H), 2.55-2.48 (m, 5H), 2.44-2.35 (m, 8H), 2.09-1.96 (m, 2H). ^13^C NMR (151 MHz, CF_3_COOD) δ 161.26, 132.70, 130.29, 127.26, 126.39, 119.28, 117.86, 52.53, 51.39, 31.89, 19.14, 17.03. HRMS-ESI, m/z calcd for C_38_H_33_Cl_4_N_4_O_4_^+^: 751.1221; found: 751.1269.

*Synthesis of PDIC-NC*: Under argon protection, the reaction mixture of 1,6,7,12-tetrachloro-3,4,9,10-tetracarboxylic dianhydride (265.0 mg, 0.5 mmol), *N,N*-dimethylethylenediamine (0.9 mL, 10 mmol), glacial acetic acid (5 mL) and NMP (5 mL) was stirred at 100 °C for 24 h. The compound N,N’-Bis(2-(dimethylammonium)ethylene)perylene-1,6,7,12-tetrachloro-3,4,9,10-tetracarboxyldiimide hydrochlorate (PDIC-NC) (222.9 mg, 60%) was then obtained as red solid after recrystallization by addition of extra HCl (2 M, 100 mL) and ethanol (100 mL) into the reaction mixture. ^1^H NMR (600 MHz, CF_3_COOD) δ 9.21 (s, 4H), 8.37 (s, 2H), 5.24 (brs, 4H), 4.22 (brs, 4H), 3.66 (s, 12H). ^13^C NMR (151 MHz, CF_3_COOD) δ 161.22, 132.72, 130.27, 127.23, 126.40, 119.21, 117.70, 54.01, 40.08, 32.30. HRMS-ESI, m/z calcd for C_32_H_25_Cl_4_N_4_O_4_^+^: 671.0595; found: 671.0551. The ^1^H NMR and ^13^C NMR data of PDIC-NC are consistent with the literature ^1^.

*Synthesis of PDI-NC*: Perylene-3,4,9,10-tetracarboxylic bisanhydride (392.1mg, 1.0 mmol), *N, N*-dimethylethylenediamine (1 mL, 9.1 mmol) were sequentially added to a reaction vial filled with deionized water under argon protection. After stirring at 100 °C for 24 h, the insoluble precipitate was filtered and washed with with 1% KOH (50 mL) and water to provide pure compound PDI-NN (425.8 mg, yield 70%) as dark purple solid. ^1^H NMR (600 MHz, CF_3_COOD) δ 9.24 (dd, *J* = 45.1, 8.1 Hz, 8H), 5.21-5.15 (m, 4H), 4.19-4.13 (m, 4H), 3.59 (s, 12H). ^13^C NMR (151 MHz, CF_3_COOD) δ 162.42, 132.61, 129.34, 125.47, 122.58, 120.60, 117.58, 54.38, 40.01, 32.34.

The compound PDI-NN was dissolved in ethanol solution of hydrochloric acid (2 M), and the reaction mixture was stirred at room temperature for another 24 h. After recrystallization in diethyl ether, N,N’-Bis(2-(dimethylammonium) ethylene) perylene-3,4,9,10- tetracarboxyldiimide hydrochlorate (PDI-NC) (537.8 mg, 85%) was produced as an orange solid. ^1^H NMR (600 MHz, CF_3_COOD) δ 9.28-9.11 (m, 8H), 8.34 (s, 2H), 5.20 (t, *J* = 4.9 Hz, 4H), 4.19 (brs, 4H), 3.61 (s, 12H). ^13^C NMR (151 MHz, CF_3_COOD) δ 162.17, 132.13, 129.07, 125.29, 122.26, 120.43, 117.79, 53.72 (d, *J* = 11.0 Hz), 40.04 (d, *J* = 13.3 Hz), 32.36 (d, *J* = 3.7 Hz). ^13^C NMR fragmentation may be caused by the formation of hydrogen bond between the hydrogen atom on the quaternary amine and the neighboring carbonyl group. HRMS-ESI, m/z calcd for C_32_H_29_N_4_O_4_^+^: 533.2184; found: 533.2186. The ^1^H NMR and ^13^C NMR data of PDI-NC are consistent with the literature ^1^.

**2 Cell culture**

MCF-7 (human breast cancer), MDA-MB-231 (human breast cancer), and HEK-293T (human embryonic kidney) cell lines were obtained from the China Center for Type Culture Collection (CCTCC). HCT-116 (human colon cancer) cells were a kind gift from Prof. Shunichi Takeda (Shenzhen University). All cell lines were maintained in high-glucose DMEM medium (Procell Life Science & Technology, Wuhan, China) supplemented with 10% fetal bovine serum (FBS; PAN-Biotech GmbH, Aidenbach, Germany) and 1% penicillin-streptomycin solution (Beyotime Biotechnology, Shanghai, China). The cells were cultured in a humidified incubator at 37 °C with 5% CO₂. Primary mouse lymphocytes were isolated from the spleen and lymph nodes of BALB/c mice. The isolated total lymphocytes were cultured in RPMI-1640 medium supplemented with 10% FBS and 1% penicillin-streptomycin for use in cytotoxicity assays.

**3** **Isolation of crude mitochondrial fraction from Cells**

Crude mitochondrial fractions were isolated using a commercial mitochondrial isolation kit (Beyotime Biotechnology, Shanghai, China) according to the manufacturer's protocol with slight modifications. Briefly, when cell confluence reached approximately 90%, monolayers were rinsed once with pre-chilled 1× PBS buffer and then detached by incubation with Trypsin-EDTA solution (Biosharp, Hefei, China). The collected cells (approx. 2 × 10⁷) were washed with PBS and then resuspended in 1 mL of ice-cold mitochondrial isolation reagent containing protease inhibitors, followed by a 15-min incubation on ice. The cell suspension was then transferred to a pre-chilled glass Dounce homogenizer (Anxiang Biochemical, Beijing, China) and homogenized with 15–20 gentle strokes on ice. The extent of cell disruption was monitored by mixing an aliquot of the homogenate with trypan blue and assessing the percentage of stained cells under a light microscope (Micro-vision, Suzhou, China); homogenization was stopped when over 80% of cells were disrupted. The homogenate was then centrifuged at 600 × g for 10 min at 4 °C to pellet nuclei and unbroken cells. The supernatant was carefully transferred to a fresh tube and centrifuged at 11,000 × g for 10 min at 4 °C to pellet the mitochondria. Isolated mitochondrial pellets were resuspended in mitochondrial storage buffer (pH7.2, Cat No. C3609, Beyotime Biotechnology, Shanghai, China), which contains ATP and sucrose to preserve mitochondria membrane potential (ΔΨm)^2^. Fluorescence-based uptake assays were conducted using an F-4600 spectrofluorometer (Hitachi, Japan) in a measurement buffer containing 10 mM HEPES (pH 6.8) and 100 mM KCl. The intrinsic fluorescence of PDIC-BL (10 µM) was first recorded. Then, the energized mitochondrial suspension was added to a final concentration of 1.25 mg/mL. To confirm the ΔΨm-dependence of this process, mitochondria were pre-incubated with the complex I inhibitor rotenone (1 µM or 10 µM) at 37°C for 30 minutes prior to addition to the PDIC-BL solution.

**4 Cytotoxicity assay**

Cell viability was determined using the MTT assay. Briefly, cells were harvested during their logarithmic growth phase using 0.25% Trypsin-EDTA (Biosharp, Hefei, China) and seeded into 96-well plates at a density of 5,000 cells/well. After a 24-hour incubation period to allow for cell attachment, the medium was replaced with fresh medium containing serial dilutions of the PDI compounds. Following a 48-hour treatment period, 10 µL of MTT solution (5 mg/mL in PBS; biofroxx, Germany) was added to each well, and the plates were incubated for an additional 4 hours at 37 °C. Subsequently, the supernatant was carefully removed, and 150 µL of dimethyl sulfoxide (DMSO; Aladdin, Shanghai, China) was added to each well to dissolve the formazan crystals. The absorbance at 570 nm was measured using a microplate reader. The half-maximal inhibitory concentration (IC_50_) values were calculated by fitting the dose-response data to a non-linear regression model using GraphPad Prism 10 software.

**5** **Colony formation assay**

To assess the long-term impact on cell proliferation, colony formation assays were performed. MCF-7 cells were seeded into 6-well plates at a density of 2,000 cells per well in complete medium and allowed to adhere for 24 hours. The treatment protocols varied depending on the compounds being compared. For the comparison between PDIC-BL and PDIC-BN, cells were treated with various concentrations of the compounds for 24 hours. Afterward, the drug-containing medium was removed, the cells were washed with PBS, and then cultured in fresh, drug-free complete medium for a total of 7 days. For the comparison between PDIC-BL and IMT1B, a continuous exposure protocol was used. Cells were cultured in complete medium containing various concentrations of the compounds for the entire 7-day duration. For all experiments, the complete medium was replaced every 3 days. At the end of the incubation period, the colonies were fixed with methanol for 1 hour, washed three times with PBS, and then stained with 0.1% crystal violet solution (Beyotime Biotechnology, Shanghai, China) for 30 minutes. After staining, the plates were gently washed with distilled water to remove excess dye, air-dried, and photographed. The number of colonies (defined as clusters of >50 cells) in each well was counted using ImageJ software, and the survival fraction was calculated relative to the untreated control group.

**6** **Comet assay (alkaline)**

DNA damage was assessed using the alkaline comet assay. MCF-7 cells were seeded in 6-well plates and allowed to adhere for 24 hours before being treated with the indicated compounds (PDI-NC, PDIC-BL, PDIC-BN) for another 24 hours. After treatment, cells were harvested, and approximately 1 × 10⁵ cells were mixed with low-melting-point agarose and layered onto slides pre-coated with normal-melting-point agarose. The slides were then immersed in a cold lysis buffer for 2 hours at 4°C. Following lysis, the slides were placed in an electrophoresis tank filled with alkaline electrophoresis buffer to allow for DNA unwinding for 30 minutes. Electrophoresis was then conducted at 25 V for 30 minutes. After electrophoresis, the slides were neutralized with a neutralization buffer, stained with ethidium bromide (EB; Aladdin, Shanghai, China), and imaged using a confocal microscope (AX, Nikon, Tokyo, Japan).

**7** **Live-cell confocal microscopy**

For all imaging experiments, MCF-7 cells were seeded onto glass-bottom confocal dishes and cultured for 24 hours to ensure adherence.For mitochondrial co-localization studies, live cells were stained with MitoTracker Deep Red FM (Beyotime Biotechnology, Shanghai, China) and Hoechst 33342 (Biosharp, Hefei, China) for 30 minutes at 37 °C. The staining solution was then replaced with fresh, pre-warmed complete medium. Baseline images were acquired before the addition of PDIC-BL (2 µM), and subsequent images were captured at various time points to observe its accumulation.For subcellular distribution comparison, cells were treated with either PDIC-BL (1 µM) or doxorubicin (1 µM; Meilunbio, Dalian, China) for the indicated durations (e.g., 3 hours) before imaging. All images were acquired using a Nikon confocal microscope (Nikon, Tokyo, Japan).

**8** **Detection of Intracellular Reactive Oxygen Species (ROS)**

Intracellular ROS levels were measured by both confocal microscopy and flow cytometry using the probe 2′,7′-dichlorofluorescin diacetate (DCFH-DA; Beyotime Biotechnology, Shanghai, China). For confocal microscopy assay, MCF-7 cells were seeded onto glass-bottom confocal dishes and allowed to adhere for 24 hours. After treatment with the indicated compounds for 6 hours, the cells were washed with serum-free medium and then incubated with 10 µM DCFH-DA at 37 °C for 20 minutes. Following incubation, the cells were washed again to remove excess probe, and fresh medium was added. The fluorescence signal was immediately captured using a Nikon confocal microscope.

For flow cytometry assay, MCF-7 cells were seeded in 6-well plates and cultured for 24 hours. After treatment with the indicated compounds for 12 hours, the cells were harvested and washed with serum-free medium. The cells were then incubated with 10 µM DCFH-DA at 37 °C for 20 minutes in the dark. After washing to remove the excess probe, the cells were resuspended in PBS and analyzed immediately on a flow cytometer (BD Biosciences, San Jose, CA, USA). Data were analyzed using FlowJo software.

For a plate-based overview of ROS production, MCF-7 cells were seeded in 6-well plates and treated with various concentrations of PDIC-BL for 48 hours. Following treatment, the cells were washed and incubated with 10 µM DCFH-DA at 37 °C for 20 minutes. The plates were then scanned using an Amersham Typhoon RGB laser-scanning imager (Cytiva, Japan). The fluorescence intensity corresponding to oxidized DCF was captured using the appropriate laser and filter settings for FITC (excitation ~488 nm, emission ~525 nm).

**9** **Immunofluorescence staining for γ-H2AX**

To detect DNA double-strand breaks, immunofluorescence staining for phosphorylated histone H2A.X (γ-H2AX) was performed. MCF-7 cells were seeded onto glass coverslips placed in 6-well plates and cultured to 70-80% confluence. The cells were then treated with the indicated compounds for 6 hours. Following treatment, the cells were washed with PBS and fixed with 4% paraformaldehyde (Servicebio, Wuhan, China) for 15 minutes at room temperature. After fixation, cells were permeabilized with 0.1% Triton X-100 in PBS for 10 minutes and then blocked with 5% bovine serum albumin (BSA) in PBS for 1 hour. The cells were subsequently incubated with a primary antibody against γ-H2AX (p-Histone H2A.X Ser139; 1:500 dilution; Beyotime Biotechnology, Shanghai, China) overnight at 4°C. The next day, after washing, the cells were incubated with a FITC-conjugated secondary antibody (1:1000 dilution; ZSGB-BIO, Beijing, China) for 1 hour at room temperature in the dark. Finally, the nuclei were counterstained with DAPI (1 µg/mL; Biosharp, Hefei, China) for 10 minutes. The coverslips were then mounted onto glass slides with an anti-fade mounting medium and imaged using a Nikon confocal microscope.

**10** **Quantitative real-time PCR (qRT-PCR)**

MCF-7 cells were seeded in 6-well plates and treated with the indicated compounds for 6 hours. Total RNA was extracted from the cells using TRIzol reagent (Biosharp, Hefei, China). The concentration and purity of the RNA were determined using a NanoDrop spectrophotometer (Thermo Fisher Scientific, Waltham, MA, USA). For cDNA synthesis, equal amounts of total RNA (approx. 3 µg) from each sample were reverse transcribed using the HiScript III RT SuperMix for qPCR (+gDNA wiper) kit (Vazyme Biotech, Nanjing, China) according to the manufacturer's protocol. The resulting cDNA was used as a template for qRT-PCR, which was performed using SYBR Green Master Mix (ACE Biotechnology, Changzhou, China) on a LightCycler 96 system (Roche, Basel, Switzerland). All primers used for detecting mitochondrial and nuclear-encoded genes were synthesized by Sangon Biotech (Shanghai, China), and their sequences are listed in Table S1. The expression of target genes was normalized to the housekeeping gene *B2M*. The relative gene expression levels were calculated using the 2-ΔΔCt method. All experiments were performed with three biological replicates.

**11** **Nude mouse xenograft model**

All animal experiments were conducted in compliance with the guidelines of the Institutional Animal Care and Use Committee (IACUC) of Huazhong University of Science and Technology. Female BALB/c nude mice (4–6 weeks old) were purchased from Beijing Vital River Laboratory Animal Technology Co., Ltd. (Beijing, China) and housed under specific pathogen-free (SPF) conditions. To establish the xenograft model, 1.5 × 10⁶ MCF-7 cells, resuspended in 100 µL of PBS, were subcutaneously injected into the right upper back of each mouse. When the average tumor volume reached approximately 50–70 mm³, the mice were randomly assigned into three groups (n = 5 per group): Control group (Vehicle, physiological saline); PDIC-BL group (2 mg/kg/day); Doxorubicin (DOX) group (3 mg/kg/day). Treatments were administered daily via intraperitoneal (i.p.) injection for 19 consecutive days. Tumor dimensions (length, L, and width, W) and body weights were recorded every two days. Tumor volume was calculated using the formula: V = (W² × L) / 2. At the end of the study, mice were euthanized, and the tumors were excised, photographed, and weighed for further analysis.

**12** **Hematoxylin and eosin (H&E) staining**

At the end of the experiment, nude mice were euthanized and dissected to remove major organs (heart, liver, spleen, lungs, and kidneys), which were immediately washed with physiological saline to remove residual blood and fixed in 10% neutral buffered formalin for 24 hours. The fixed tissues were dehydrated through a graded ethanol series, cleared with xylene (Servicebio, Wuhan, China), embedded in paraffin, and sectioned into continuous slices with a thickness of 4 µm. After deparaffinization with xylene and rehydration through graded ethanol, the sections were stained with hematoxylin (Servicebio, Wuhan, China) for 5 minutes, rinsed with running water, differentiated with 1% hydrochloric acid ethanol, and then blued with running water. Subsequently, the cytoplasm was stained with eosin for 2 minutes, dehydrated through graded ethanol, cleared with xylene, and coverslips were mounted using neutral resin. The sections were observed under an optical microscope for tissue morphology and subjected to pathological evaluation.

**13** **TUNEL staining**

Tumor tissues dissected from nude mice were fixed in 10% neutral buffered formalin for 24 hours, then dehydrated through a graded ethanol series, cleared with xylene, embedded in paraffin, and sectioned. Sections were subjected to antigen retrieval with proteinase K (original solution: PBS=1:9, Servicebio, Wuhan, China) at 37°C for 15 minutes, then washed with PBS three times for 5 minutes each. Appropriate amounts of TDT enzyme, dUTP, and buffer from the TUNEL kit (Servicebio, Wuhan, China) were mixed in a ratio of 1:5:50 according to the number of sections and tissue size, added to cover the tissue within the ring, and incubated in a humidified chamber at 37°C for 1 hour with a small amount of water added to maintain humidity. DAPI was used to counterstain the cell nuclei: sections were washed with PBS (pH 7.4) three times, each for 5 minutes. After removing PBS, DAPI staining solution was added within the ring, and the sections were incubated in the dark at room temperature for 10 minutes. Finally, slides were coverslipped with anti-fade mounting medium and observed under a fluorescence microscope. Red signals represent TUNEL-positive cells, and blue signals represent DAPI-stained nuclei.

**14** **Single molecule stretching assay**

Single-molecule magnetic tweezers were used for the stretching assay (BioPSI, Singapore). The measurements were conducted with a 5,901 bp PCR product spanning the mitochondrial ND1–ATP6 region, amplified from genomic DNA extracted from MCF-7 cells using the Foregene DNA Extraction Kit (Foregene, China). A pair of primers labeled with biotin and a thiol group at their respective 5′ ends (Sangon Biotech Co., Ltd., China) was used for the amplification. A flow-channel was prepared as previously described ^3^. The 5’-thiol end of dsDNA was attached to the coverslip surface via a sulfosuccinimidyl 4-(*N*-maleimidomethyl) cyclohexane-1-carboxylate (sulfo-SMCC) crosslinker (Huateng Pharma Co., Ltd, China), while the 5’-biotin-end of dsDNA was attached to 2.8 µm-diameter streptavidin-coated paramagnetic beads Dynal M280 (Thermo Fisher Scientific, USA). The force was applied to the magnetic beads by a pair of permanent magnets positioned above sample. The bead-height was measured from the diffraction patterns of the beads using magnetic tweezers. The bead-height force curves were obtained through a force-jump procedure, and at each force, the magnet was held for 5 s to measure the bead-heights. All single-molecule experiments were conducted at room temperature of 20-23°C.

**15** **LC-MS analysis**

The filtered samples were analyzed by an Agilent & 1290 Infinity II / 6545 QTOF LC/MS using an Agilent Zorbax Eclipse Plus C18 RRHD column (1.8 μm, 50 × 2.1 mm) with a linear gradient of 5% to 100% solvent A (solvent A: 0.1% HCOOH in CH_3_CN; solvent B: 0.1% HCOOH in H_2_O) over 17 min at a flow rate of 0.4 mL/min.

**16** **ROS scavenging and cytotoxicity rescue assay**

MCF-7 cells were seeded onto glass-bottom confocal dishes. After 24 hours, the cells were pre-treated with complete medium containing 5 mM N-acetylcysteine (NAC; Aladdin, Shanghai, China) for 4 hours. The medium was then replaced with fresh medium containing 10 µM PDIC-BL, with or without 5 mM NAC, and the cells were incubated for an additional 6 hours. Intracellular ROS levels were then detected by staining with 10 µM DCFH-DA for 20 minutes, followed by imaging on a Nikon AX confocal microscope (Nikon, Tokyo, Japan). MCF-7 cells were seeded in 96-well plates. After 24 hours, the cells were pre-treated with complete medium containing 5 mM NAC for 4 hours. Subsequently, the medium was replaced with fresh medium containing serial dilutions of PDIC-BL, with or without 5 mM NAC. The cells were then incubated for 24 hours. Cell viability was assessed using the standard MTT assay as described in section 4.

**Supplementary Tables**

**Table S1. IC_50_ values of compounds against MCF-7 and MDA-MB-231 breast cancer cells**

| Compound | MCF-7 IC_50_ (μM) | MDA-MB-231 IC_50_ (μM) |
| --- | --- | --- |
| PDIC-BL | 0.59 ± 0.09 | 0.48 ± 0.02 |
| PDIC-PD | 5.0 ± 1.0 | 2.2 ± 0.9 |
| PDIC-NC | 0.8 ± 0.2 | 0.39 ± 0.02 |
| PDIC-BN | > 10 | > 10 |
| PDI-NC | 3.0 ± 1.3 | 0.39 ± 0.02 |
| IMT1B | > 30 | > 30 |

**Table S2. Primers used for mitochondrial gene single-molecule stretching assay**

| Primer Name | Sequence (5′→3′) |
| --- | --- |
| mtND1 biotin Forward | AAGTCACCCTAGCCATCATTCTAC |
| mtND1 thiol Reverse | GCGACAGCGATTTCTAGGATAG |

Note: all primers are oriented 5′→3′.

**Table S3. Primers used for mitochondrial gene qPCR analysis**

| Primer Name | Sequence (5′→3′) |
| --- | --- |
| mt12S Forward | AAACTGGGATTAGATACCCC |
| mt12S Reverse | GAGGGTGACGGGCGGTGTGT |
| mt16S Forward | CGGTTTGAACTCAGATCACGT |
| mt16S Reverse | CTCCGGTCTGAACTCAGATCACGT |
| mtCOX3 Forward | ACAGGGTTTGGAGCCAACTA |
| mtCOX3 Reverse | GTGCCGATACCGTGTAGGTT |
| mtATP6 Forward | CCATCAGCCTACTCATTCAACC |
| mtATP6 Reverse | GCGACAGCGATTTCTAGGATAG |
| mtCYTB Forward | TATTCGCCTACACAATTCTCCG |
| mtCYTB Reverse | GCTTACTGGTTGTCCTCCGATT |
| mtND1 Forward | AAGTCACCCTAGCCATCATTCTAC |
| mtND1 Reverse | GCAGGAGTAATCAGAGGTGTTCTT |
| mtCO1 Forward | TACGTTGTAGCCCACTTCCACT |
| mtCO1 Reverse | GGATAGGCCGAGAAAGTGTTGT |
| mttRNA-Leu (UUR) Forward | CACCCAAGAACAGGGTTTGT |
| mttRNA-Leu (UUR) Reverse | TGGCCATGGGTATGTTGTTA |
| mtCOII Forward | CCCCACATTAGGCTTAAAAACAGAT |
| mtCOII Reverse | TATACCCCCGGTCGTGTAGCGGT |
| B2M Forward | TGCTGTCTCCATGTTTGATGTATCT |
| B2M Reverse | TCTCTGCTCCCCACCTCTAAGT |

Note: all primers are oriented 5′→3′.

**Supplementary Figures**

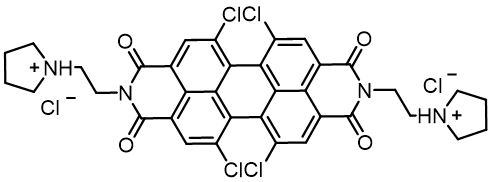

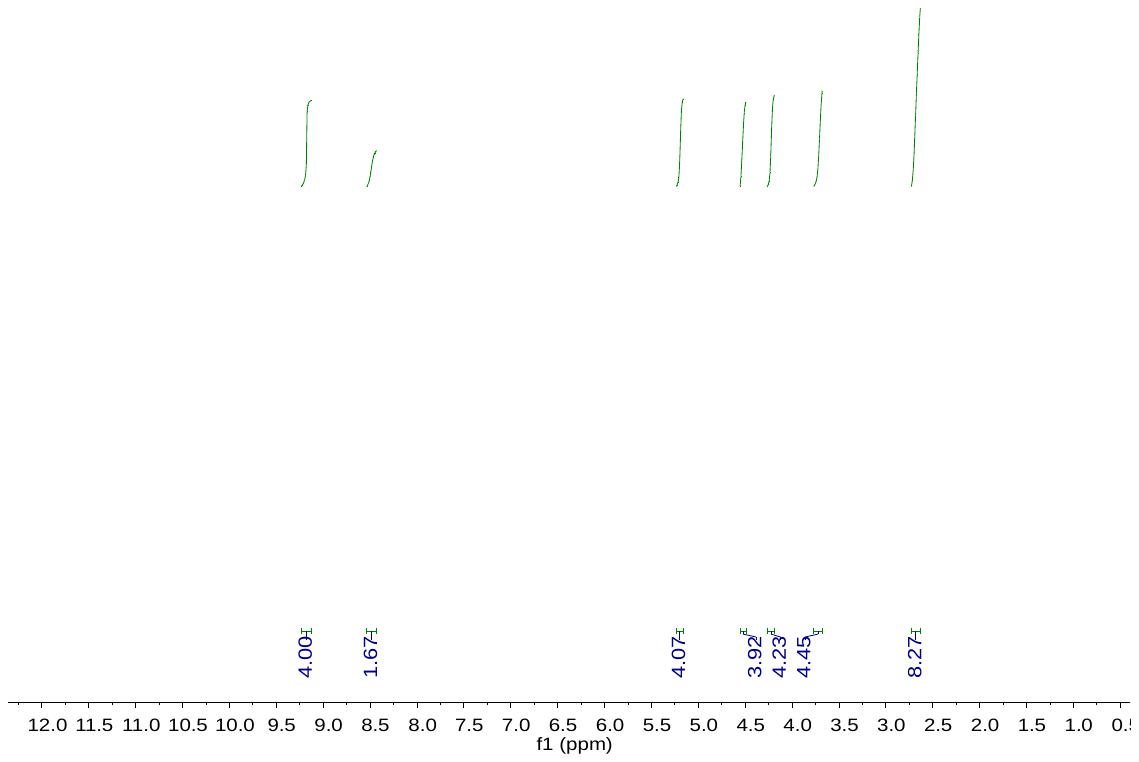

**Figure S1**. The ^1^H NMR of PDIC-BL.

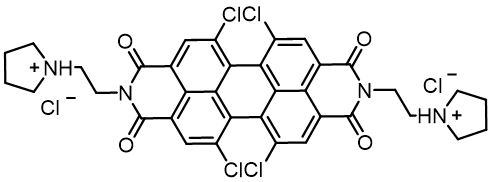

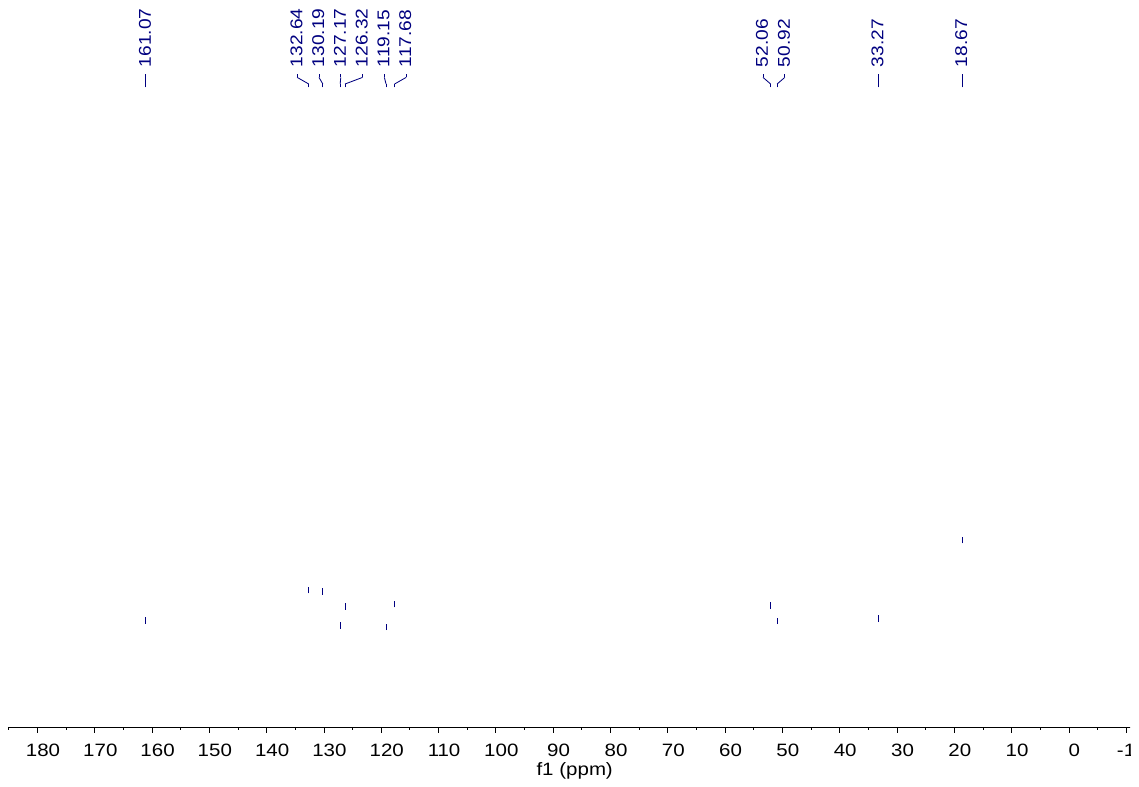

**Figure S2**. The ^13^C NMR of PDIC-BL.

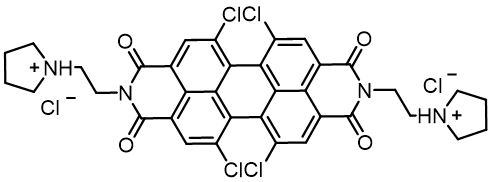

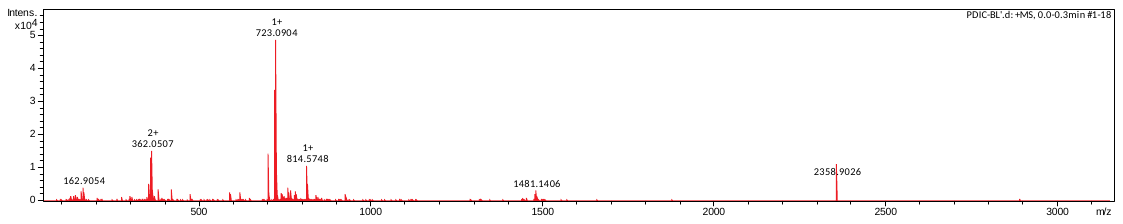

**Figure S3**. The HRMS-ESI of PDIC-BL.

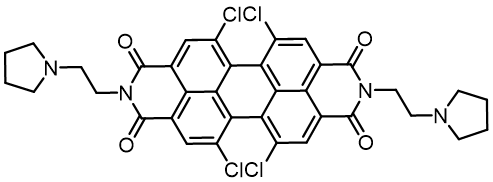

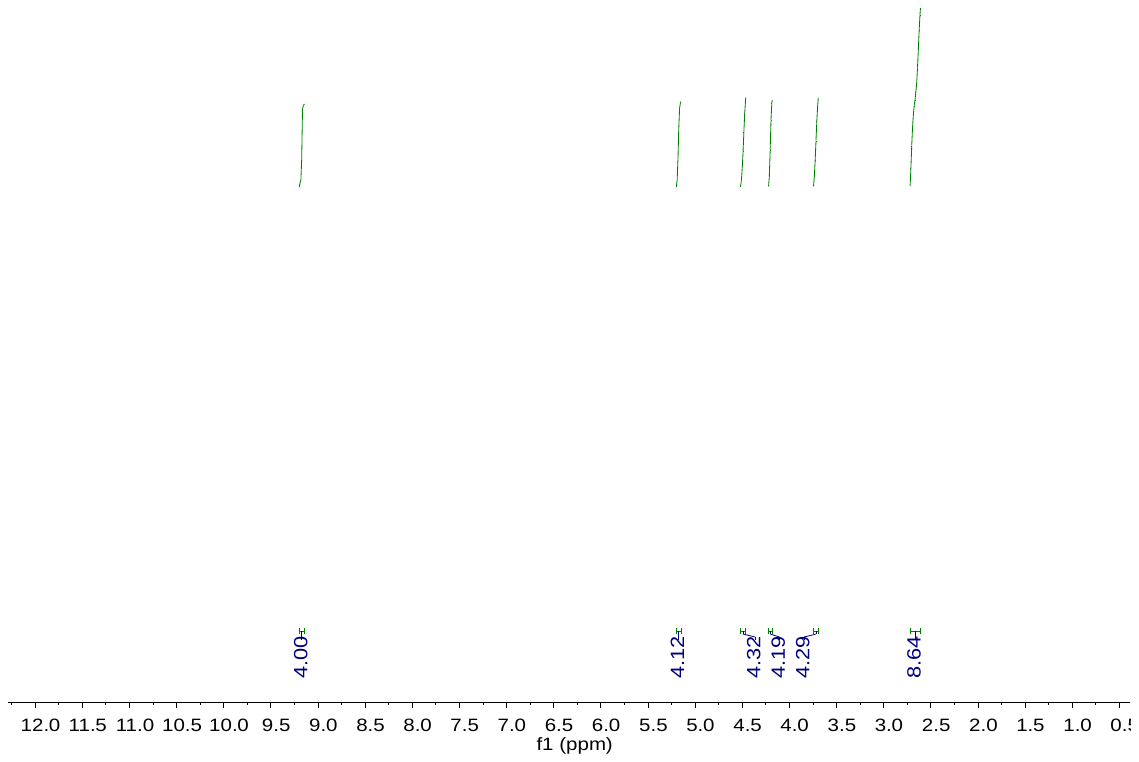

**Figure S4.** The ^1^H NMR of PDIC-BN.

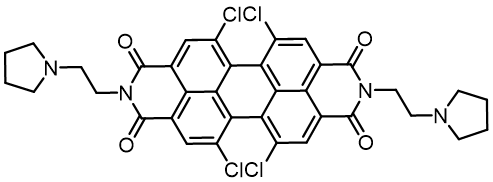

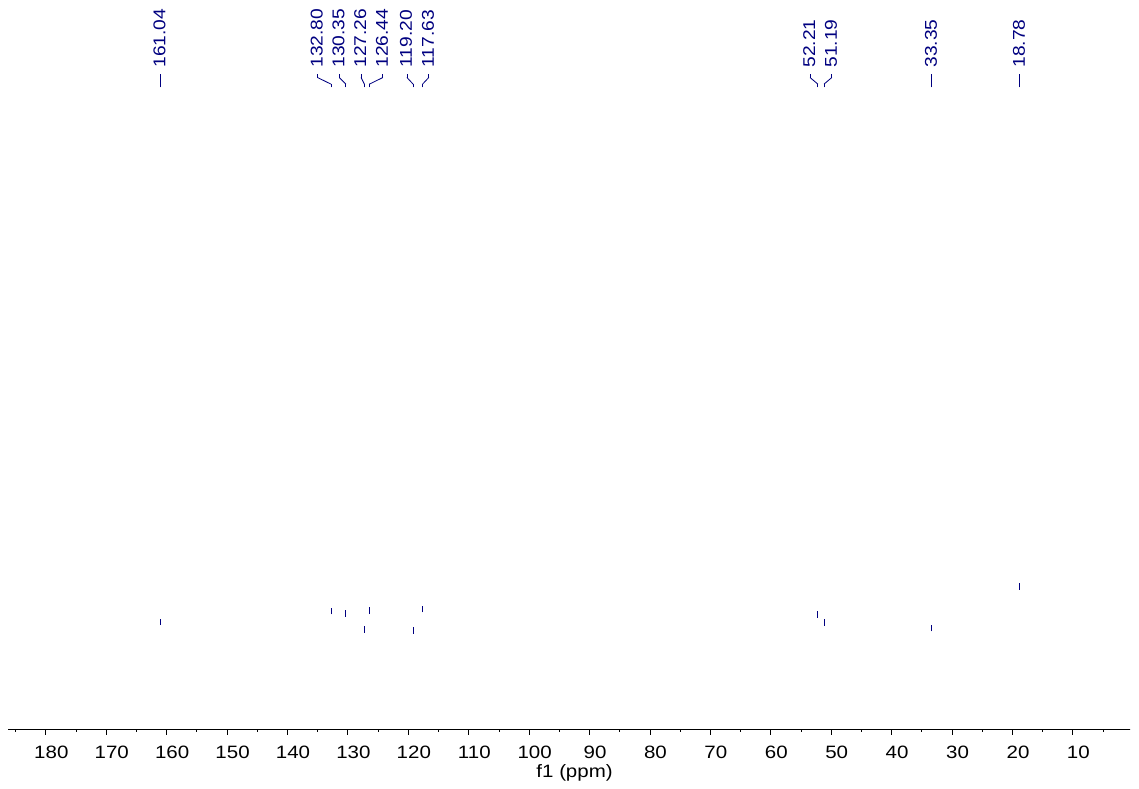

**Figure S5.** The ^13^C NMR of PDIC-BN.

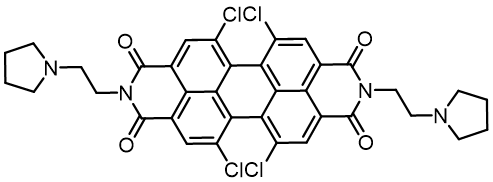
**Figure S6.** The
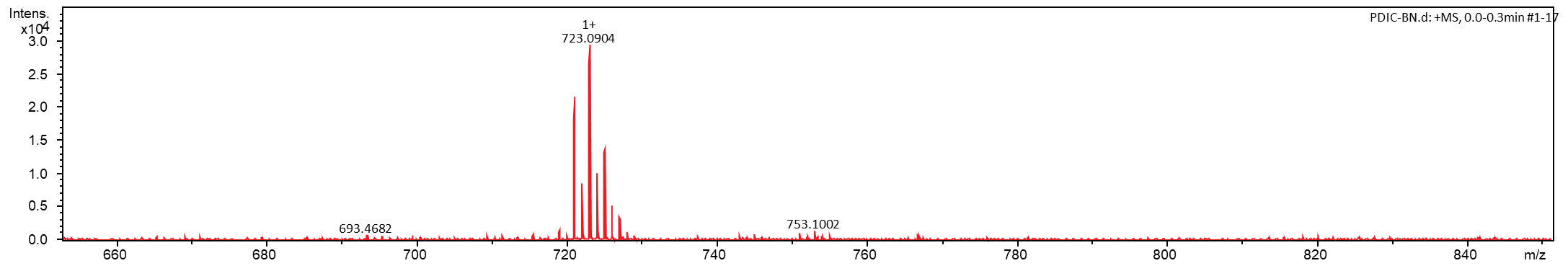
HRMS-ESI of PDIC-BN.

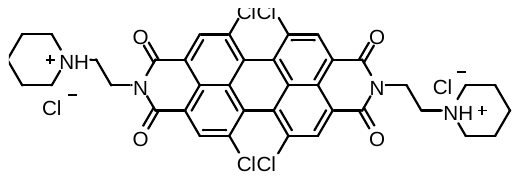

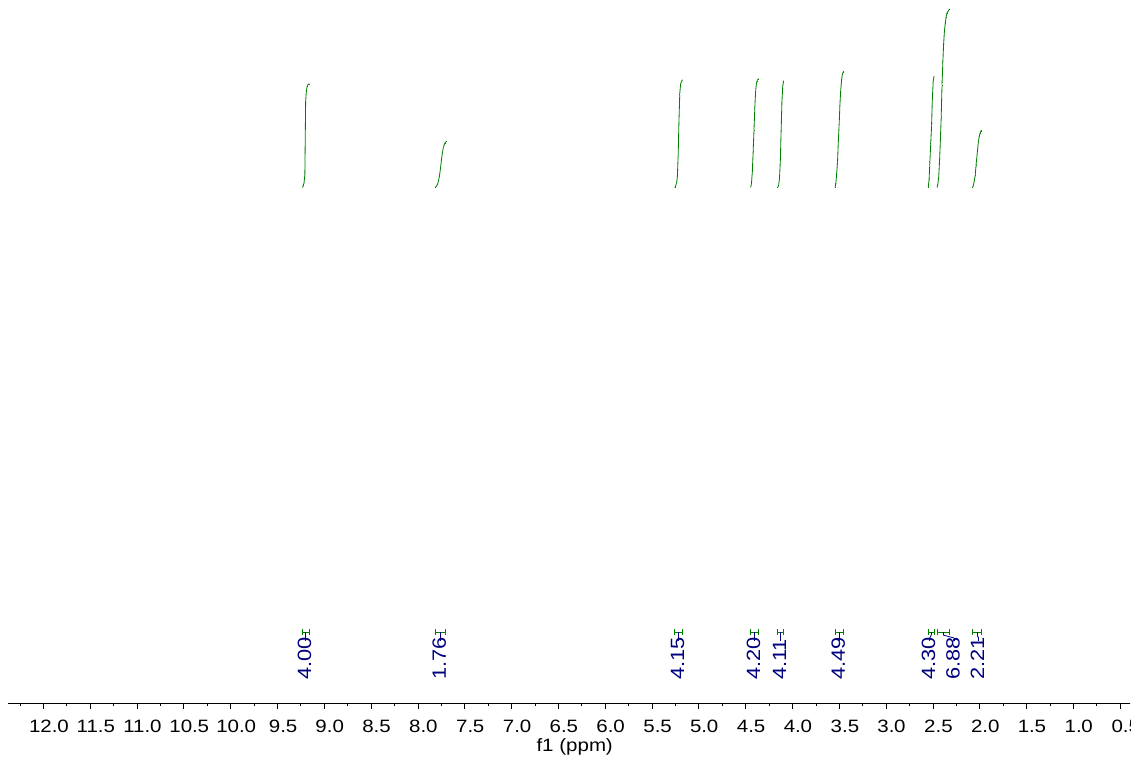

**Figure S7.** The ^1^H NMR of PDIC-PD.

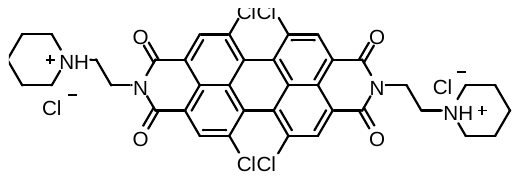

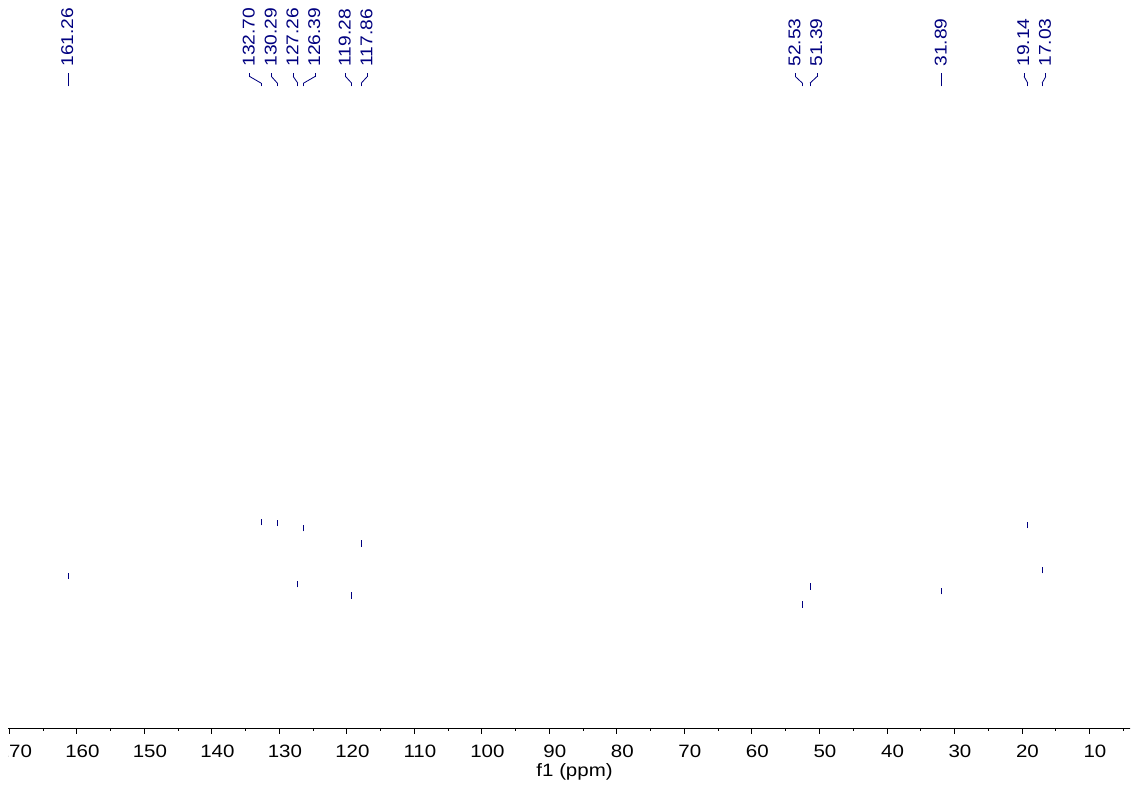

**Figure S8.** The ^13^C NMR of PDIC-PD.

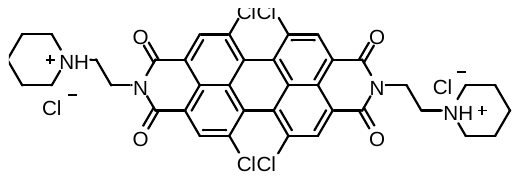

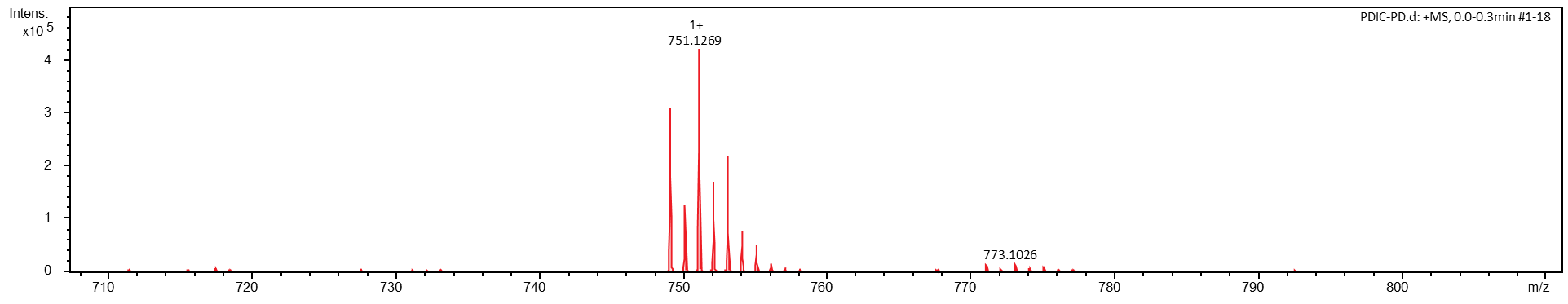

**Figure S9.** The HRMS-ESI of PDIC-PD.

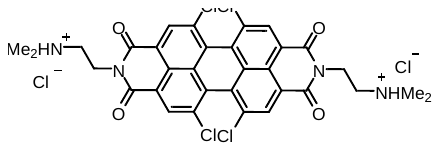

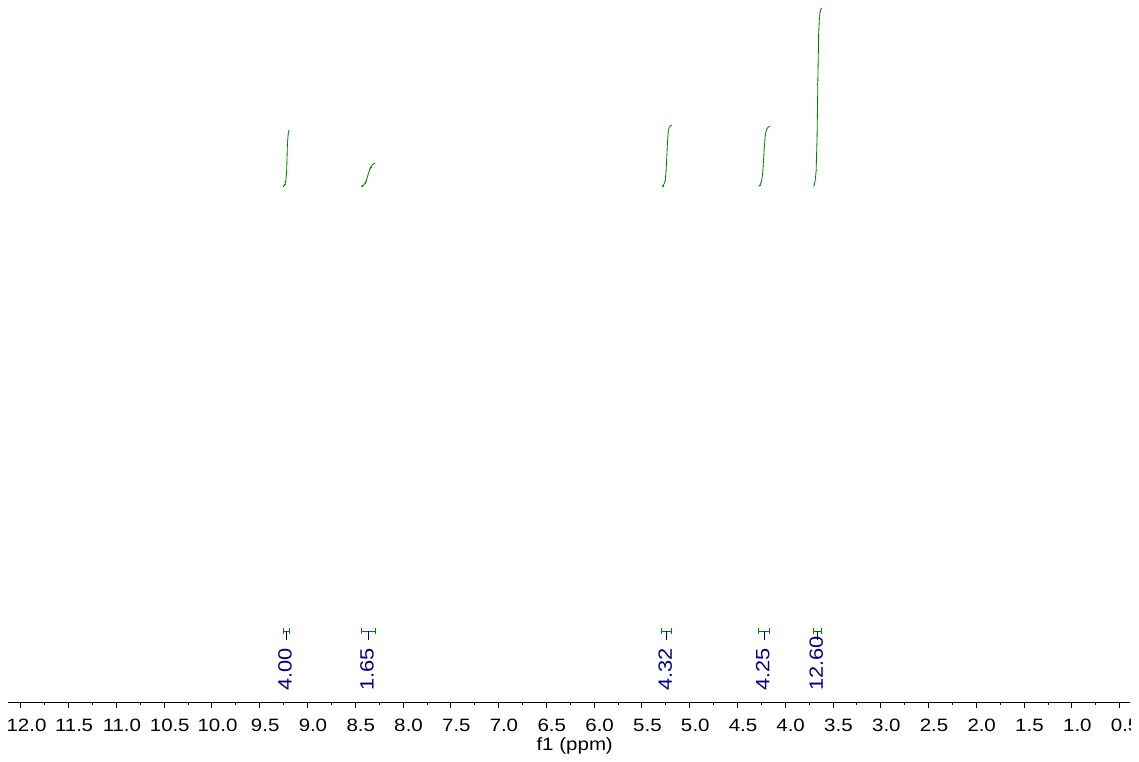

**Figure S10**. The ^1^H NMR of PDIC-NC.

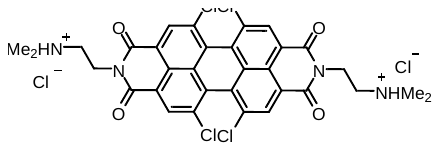

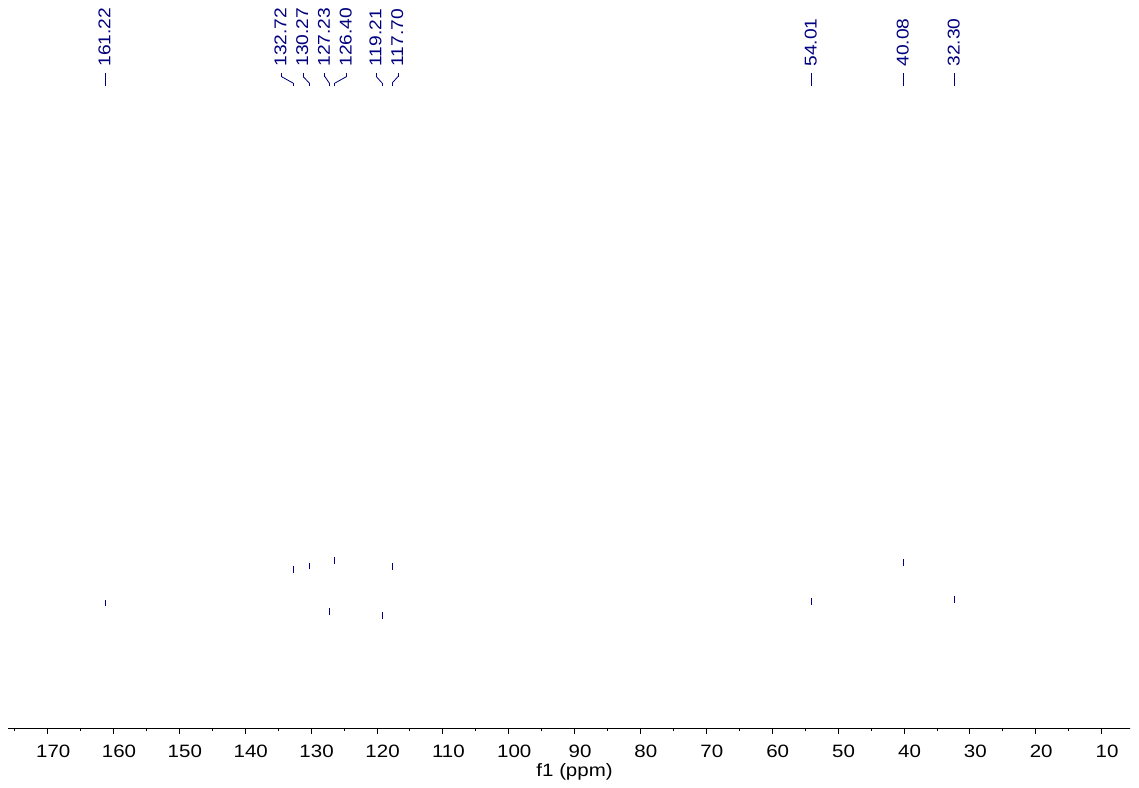

**Figure S11**. The ^13^C NMR of PDIC-NC.

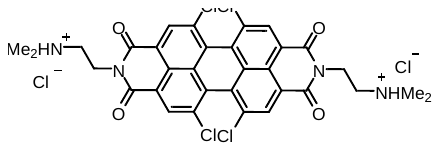

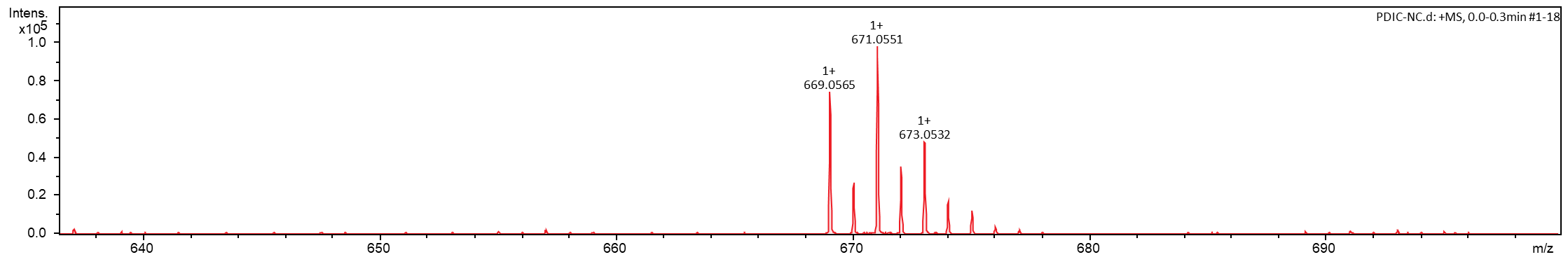

**Figure S12**. The HRMS-ESI of PDIC-NC.

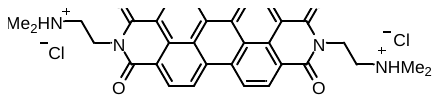

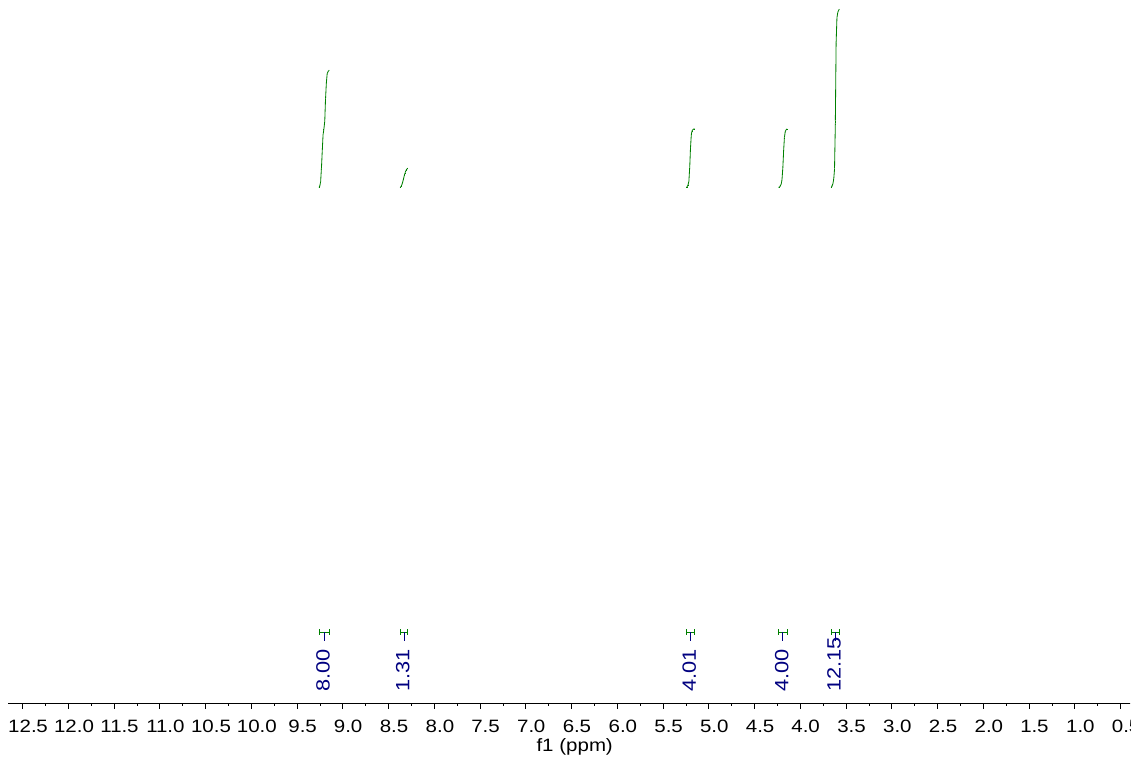

**Figure S13**. The ^1^H NMR of PDI-NC.

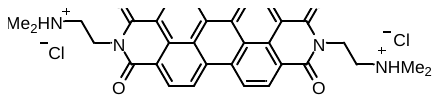

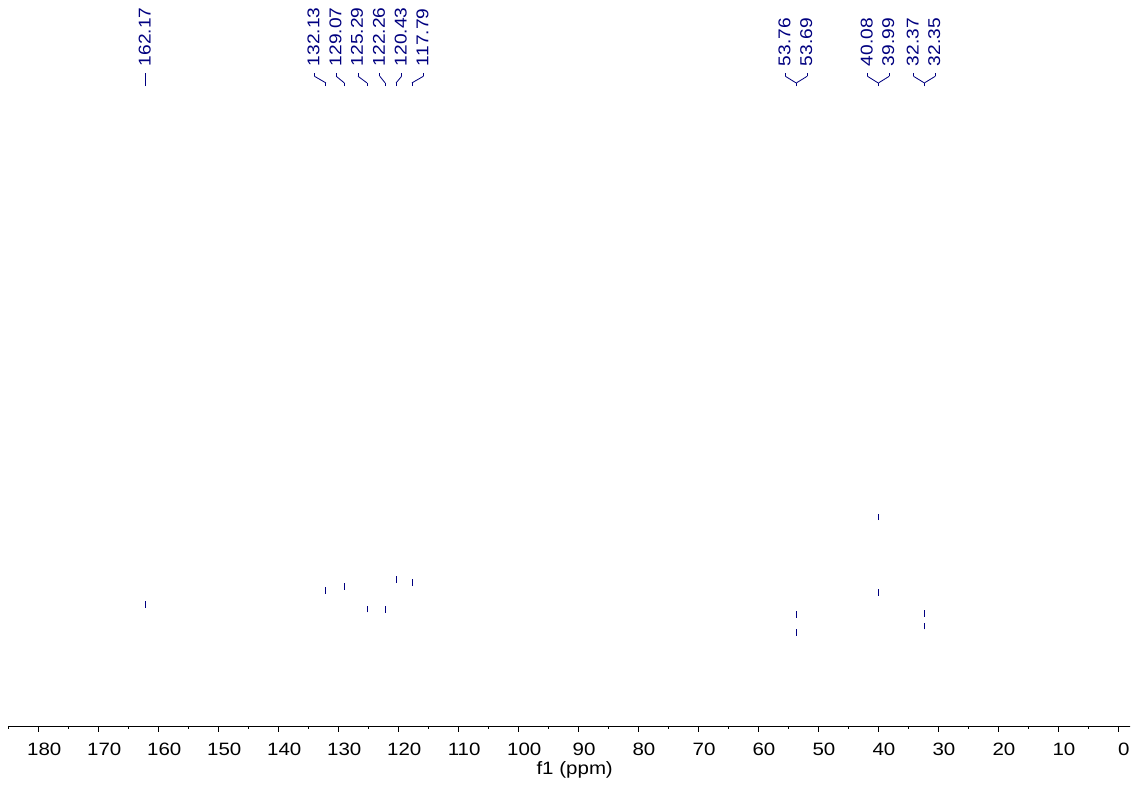

**Figure S14**. The ^13^C NMR of PDI-NC.

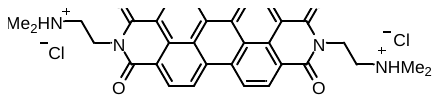

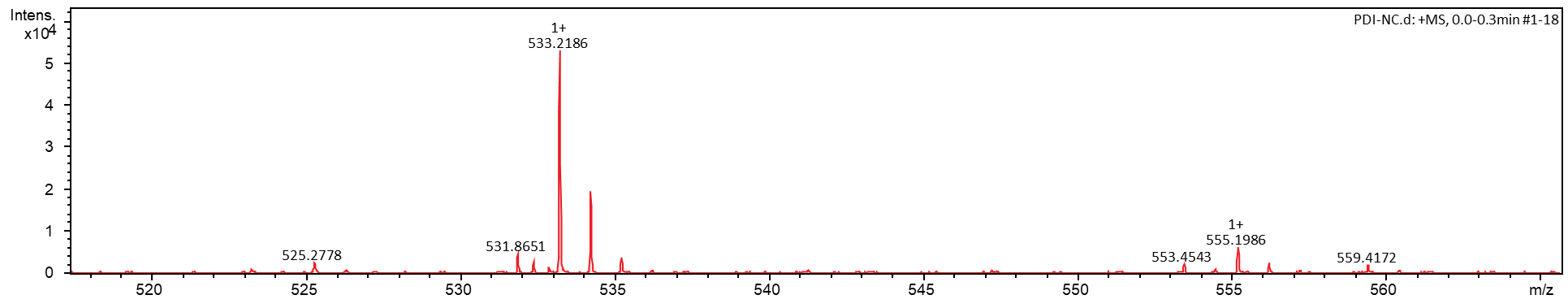

**Figure S15**. The HRMS-ESI of PDI-NC.

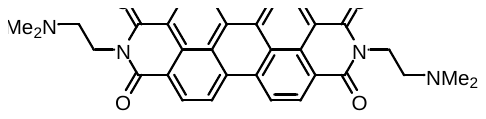

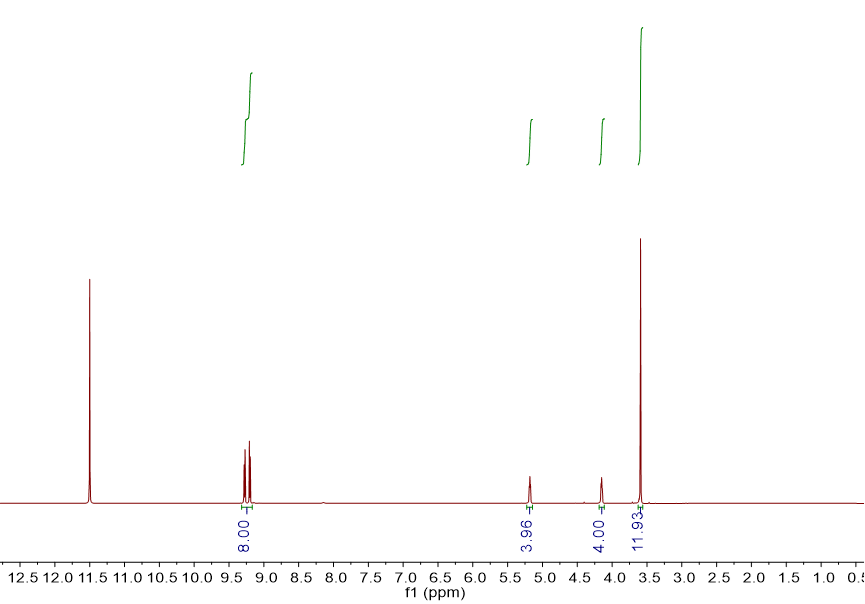

**Figure S16**: The ^1^H NMR of PDI-NN.

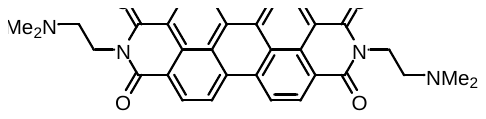

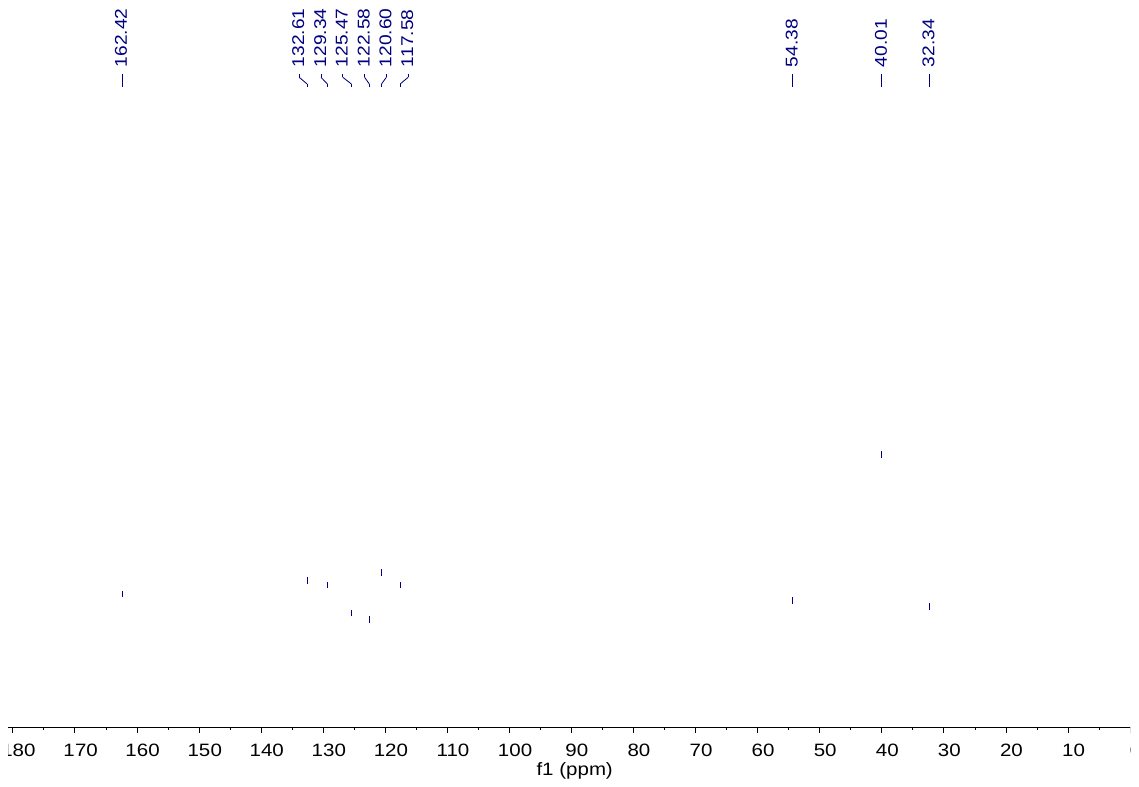

**Figure S17**. The ^13^C NMR of PDI-NN.

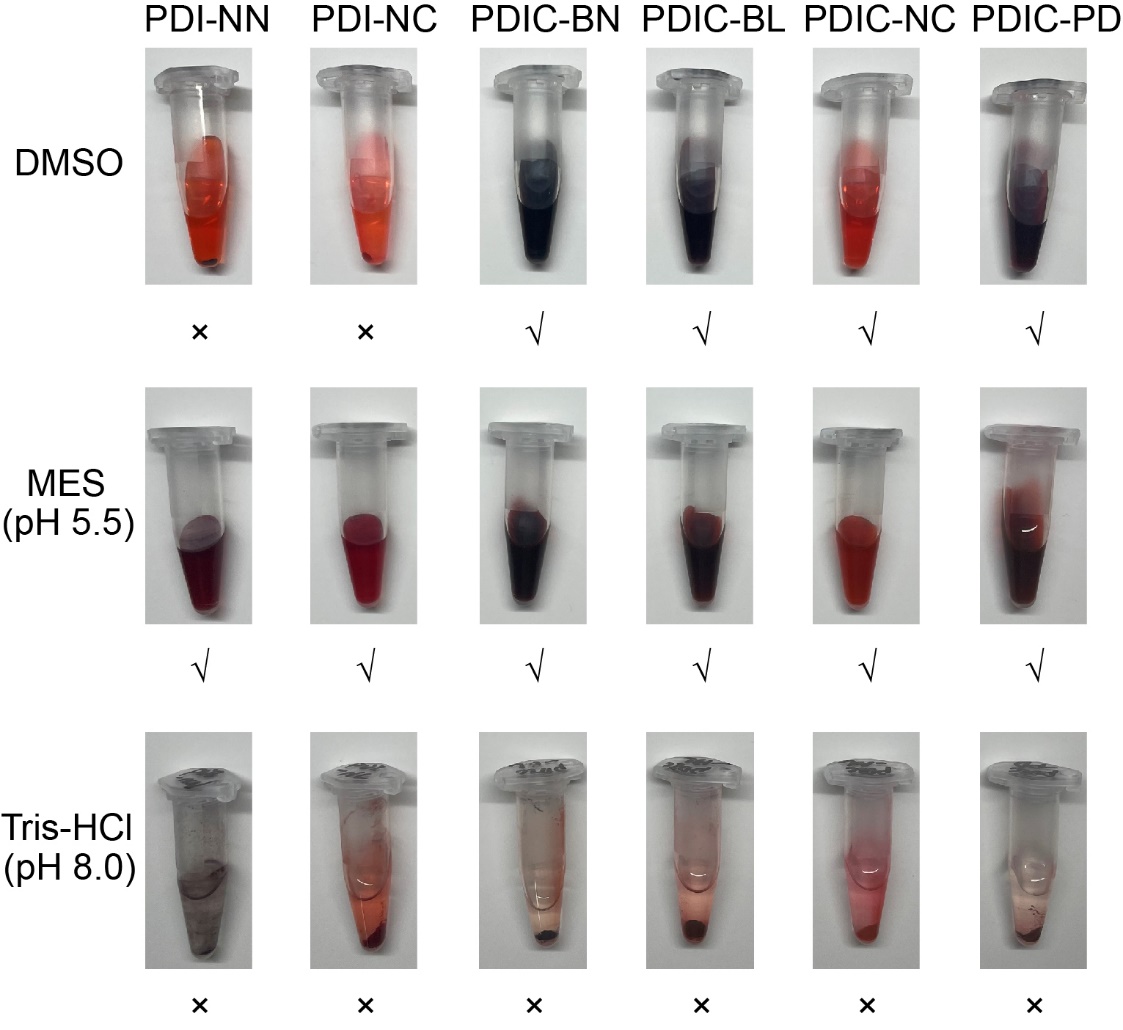

**Figure S18. Solubility profiles of PDI derivatives under different solvent conditions.** Three solvent condition was used, including dimethyl sulfoxide (DMSO), MES buffer (pH 5.5), and Tris-HCl buffer (pH 8.0). Complete dissolution is indicated by "√", while partial dissolution or no dissolution (e.g., visible precipitation) is marked with "×". The results demonstrate that bay-region chlorination (PDIC series) significantly enhances the solubility in DMSO. Additionally, acidic condition (MES buffer) facilitates the dissolution of all compounds, suggesting a pH-dependent solubility profile of these derivatives.

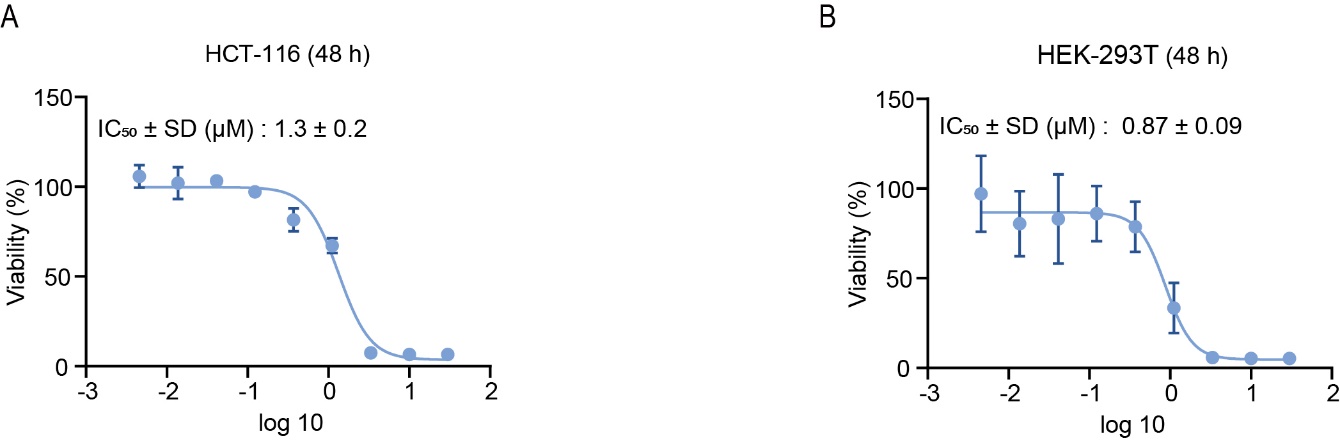

**Figure S19. Cytotoxicity of PDIC-BL in various cancer cell lines.** (**A**-**B**) Dose-response curves showing the cytotoxic effect of PDIC-BL on (A) HCT-116 and (B) HEK-293T cells after 48 hours of treatment. Cell viability was determined using the MTT assay. The calculated IC_50_ values were 1.3 ± 0.2 µM for HCT-116 cells and 0.87 ± 0.09 µM for HEK-293T. Data are presented as mean ± SD (n = 3).

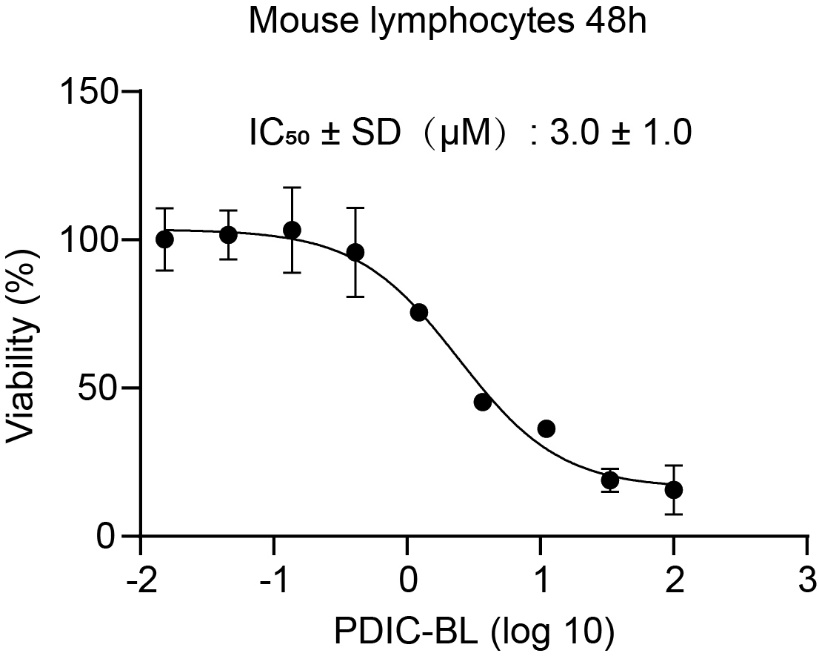

**Figure S20.** **Cytotoxicity of PDIC-BL in mouse lymphocyte cells**. Mouse lymphocytes were incubated with PDIC-BL at concentrations ranging from 100 µM down to ~0.015 µM (three-fold serial dilutions) for 48 h. Cell survival was determined by MTT assay and expressed as a percentage of untreated control. Data points represent mean ± SD (n = 3), and the solid line is a nonlinear fit to a sigmoidal dose–response model plotted against the log_10_ of PDIC-BL concentration. The calculated IC_50_ ± SD is 3.0 ± 1.0 µM.

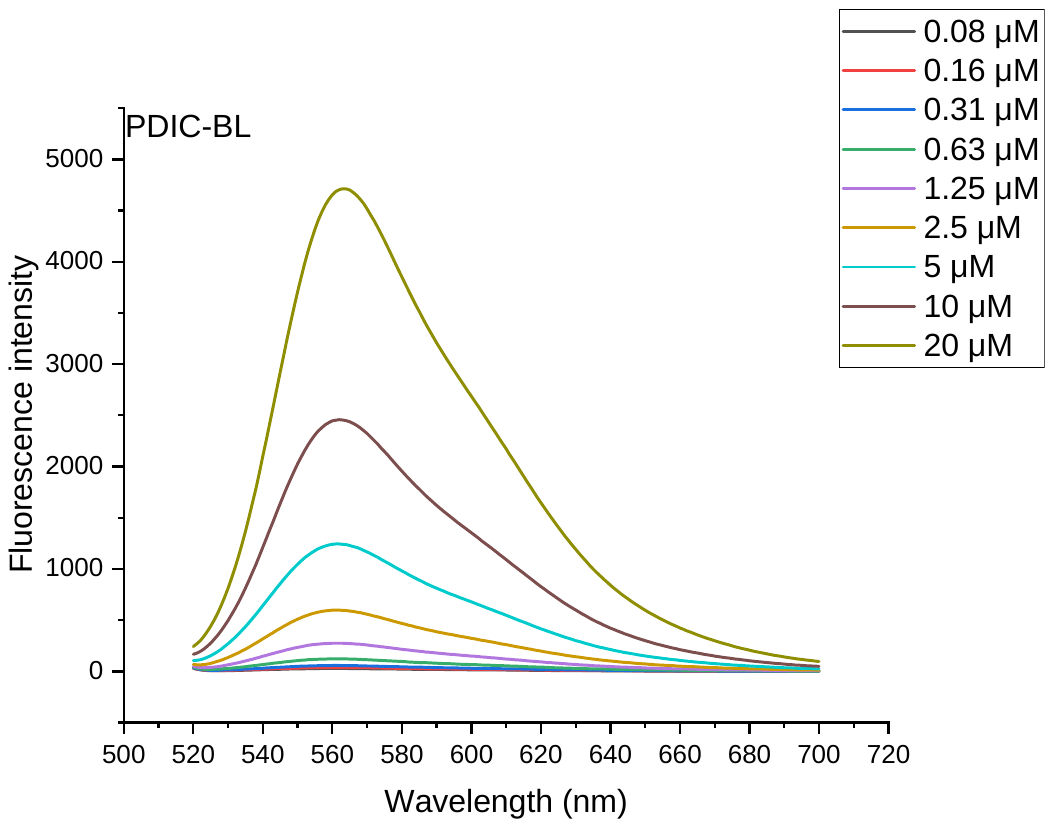

**Figure S21. Concentration-dependent fluorescence emission of PDIC-BL**. (A) PDIC-BL was prepared at varying concentrations (0.08, 0.16, 0.31, 0.63, 1.25, 2.5, 5, 10, and 20 µM) in the same buffer. Samples were excited at 488 nm, and emission spectra were recorded from 500 to 720 nm under identical instrumental settings. Emission intensity increases monotonically with PDIC-BL concentration, with a maximum at 565 nm.

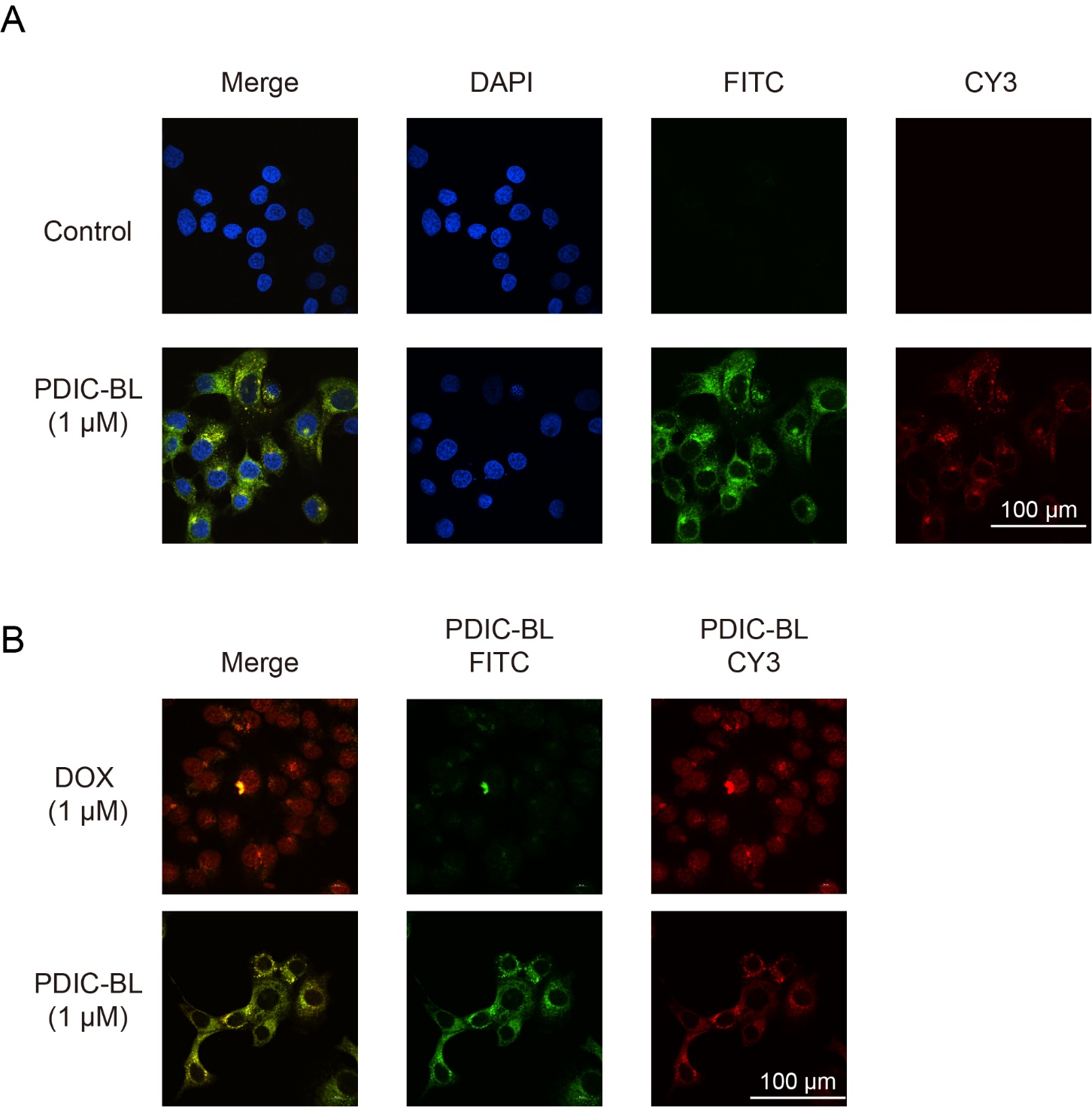

**Figure S22.** **Confocal imaging of PDIC-BL cellular uptake and comparison with doxorubicin.** (A) Cells were treated with 1 µM PDIC-BL or vehicle (Control) for 24 h and co-stained with DAPI to label nuclei (blue). PDIC-BL auto-fluoresces in both the green and red channels, showing its intracellular distribution in live cells. (B) Cells were treated for 3 h with 1 µM doxorubicin (DOX) or PDIC-BL. DOX (red channel) localizes predominantly to the nucleus, whereas PDIC-BL (green/red channels) accumulates throughout the cytoplasm, consistent with mitochondrial targeting.

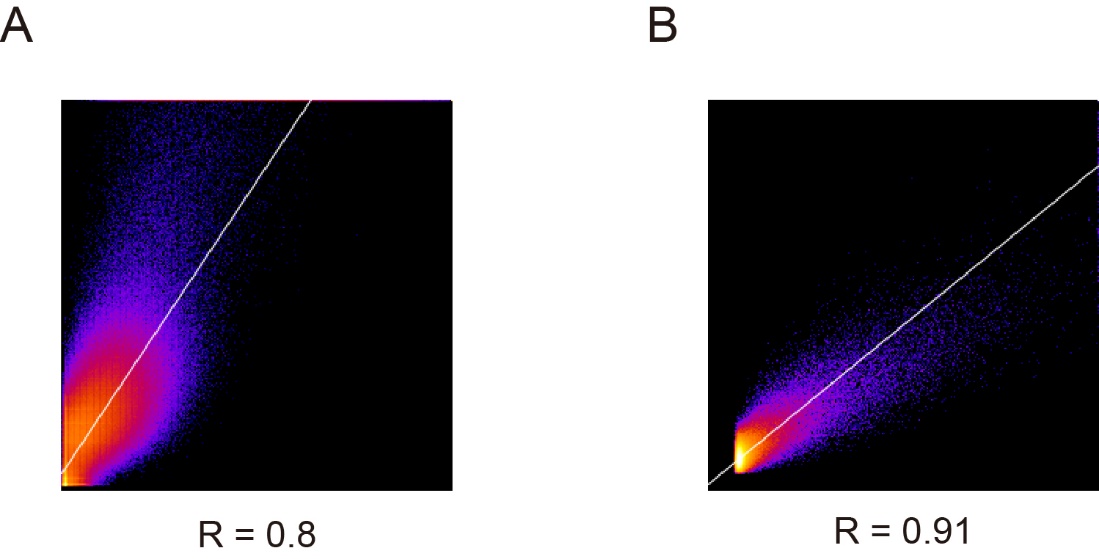

**Figure S23. Quantitative colocalization analysis of PDIC-BL and mitochondria**. Quantitative analysis was performed using the Coloc 2 plugin in ImageJ (https://imagej.net/imaging/colocalization-analysis). (A) 2D intensity histogram for PDIC-BL (green channel) and MitoTracker (far-red channel) signals in MCF-7 cells, corresponding to main text Figure 2C. The analysis yielded a Pearson's correlation coefficient of 0.80. (B) 2D intensity histogram for PDIC-BL and MitoTracker signals in isolated mitochondria, corresponding to main text Figure 2E. The analysis yielded a Pearson's correlation coefficient of 0.91.

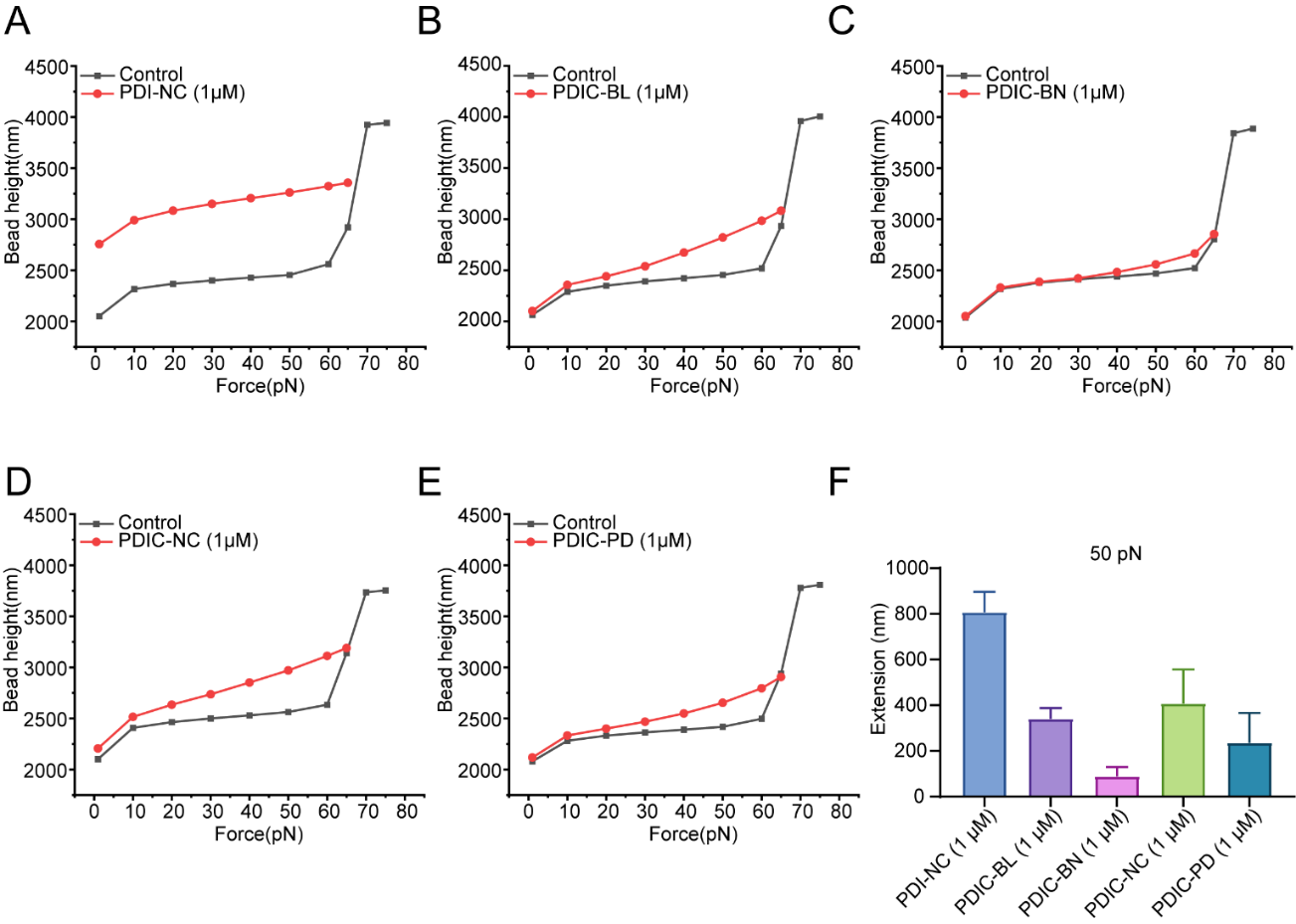

**Figure S24.** **DNA binding properties of PDIs analyzed using single-molecule stretching assay.** (A-E) Mean force-extension curves (n=3) of a single dsDNA molecule upon addition of 1 µM of the indicated PDI derivative. The control curve (no compound) is shown for comparison. (A) PDI-NC, (B) PDIC-BL, (C) PDIC-BN, (D) PDIC-NC, and (E) PDIC-PD. (F) Quantitative comparison of the DNA extension (in nm) induced by each compound (1 µM) at 50 pN, with data extracted from the force-extension curves. The results demonstrate that the planar PDI-NC is the most potent intercalator. The non-planar, cationic derivatives (PDIC-BL, PDIC-NC, PDIC-PD) retain significant, albeit varied, intercalation activity. In contrast, the non-planar, neutral analogue PDIC-BN shows minimal interaction with DNA, underscoring the critical role of the cationic side chains for DNA binding. Data in (F) are presented as mean ± SD (n = 3).

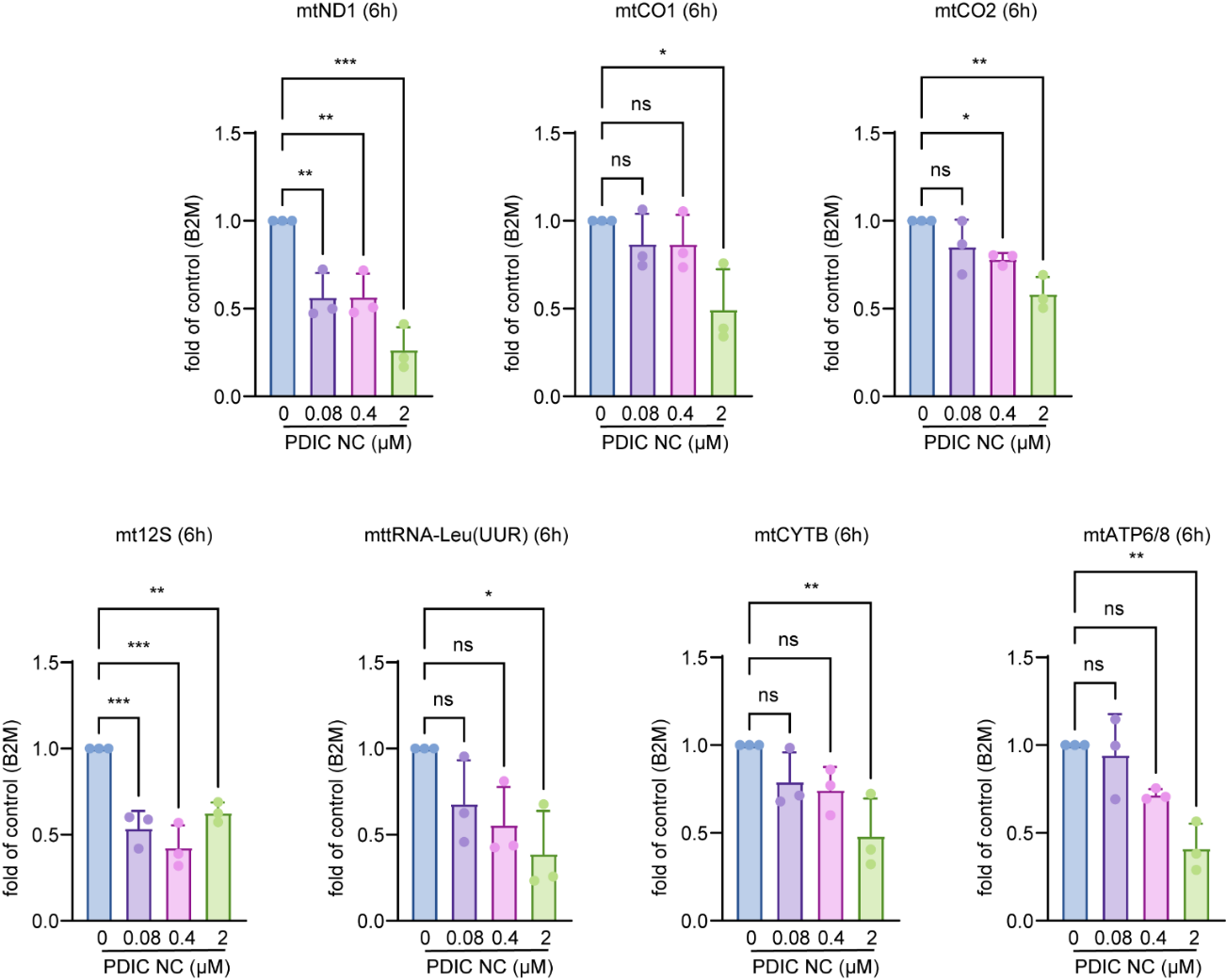

**Figure S25.PDIC-NC suppress mtDNA transcription in MCF-7 cells.** MCF-7 cells were treated with PDIC-NC at 0, 0.08, 0.4, or 2 µM for 6 h. Total RNA was extracted, and transcript levels of mtND1, mtCO1, mtCO2, mt12S, mttRNA-Leu (UUR), mtCYTB and mtATP6/8 were quantified by qRT-PCR using B2M as an internal control. Data are presented as fold change relative to untreated control (mean ± SD, n = 3). Statistical comparisons were made against the 0 µM group: **p* < 0.05, ***p* < 0.01, ****p* < 0.001,; ns, not significant.

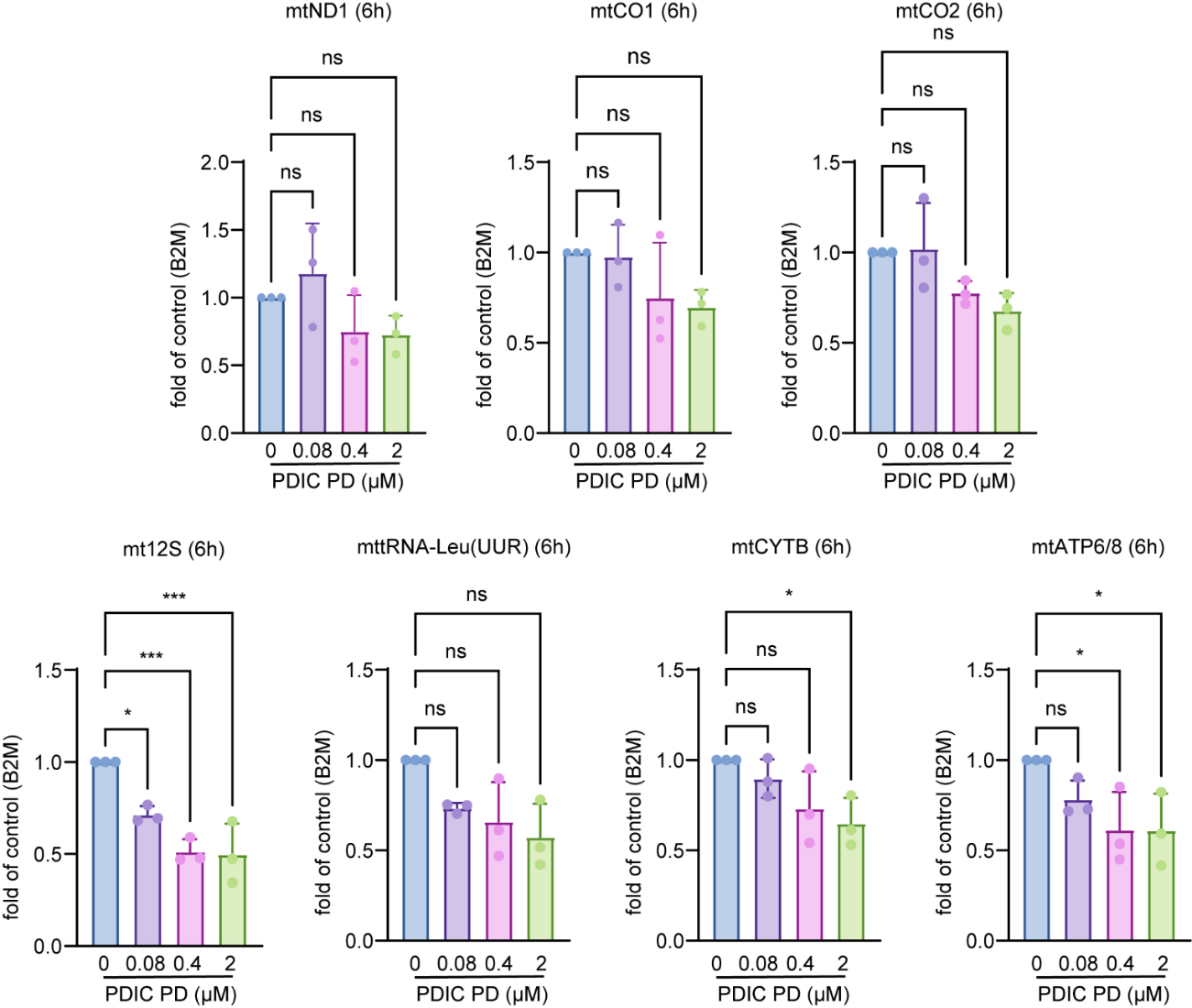

**Figure S26.** **PDIC-PD suppress mtDNA transcription in MCF-7 cells.** MCF-7 cells were treated with PDIC-PD at 0, 0.08, 0.4, or 2 µM for 6 h. Total RNA was extracted, and transcript levels of mtND1, mtCO1, mtCO2, mt12S, mttRNA-Leu (UUR), mtCYTB and mtATP6/8 were quantified by qRT-PCR using B2M as an internal control. Data are presented as fold change relative to untreated control (mean ± SD, n = 3). Statistical comparisons were made against the 0 µM group: **p* < 0.05, ***p* < 0.01, ****p* < 0.001, ; ns, not significant.

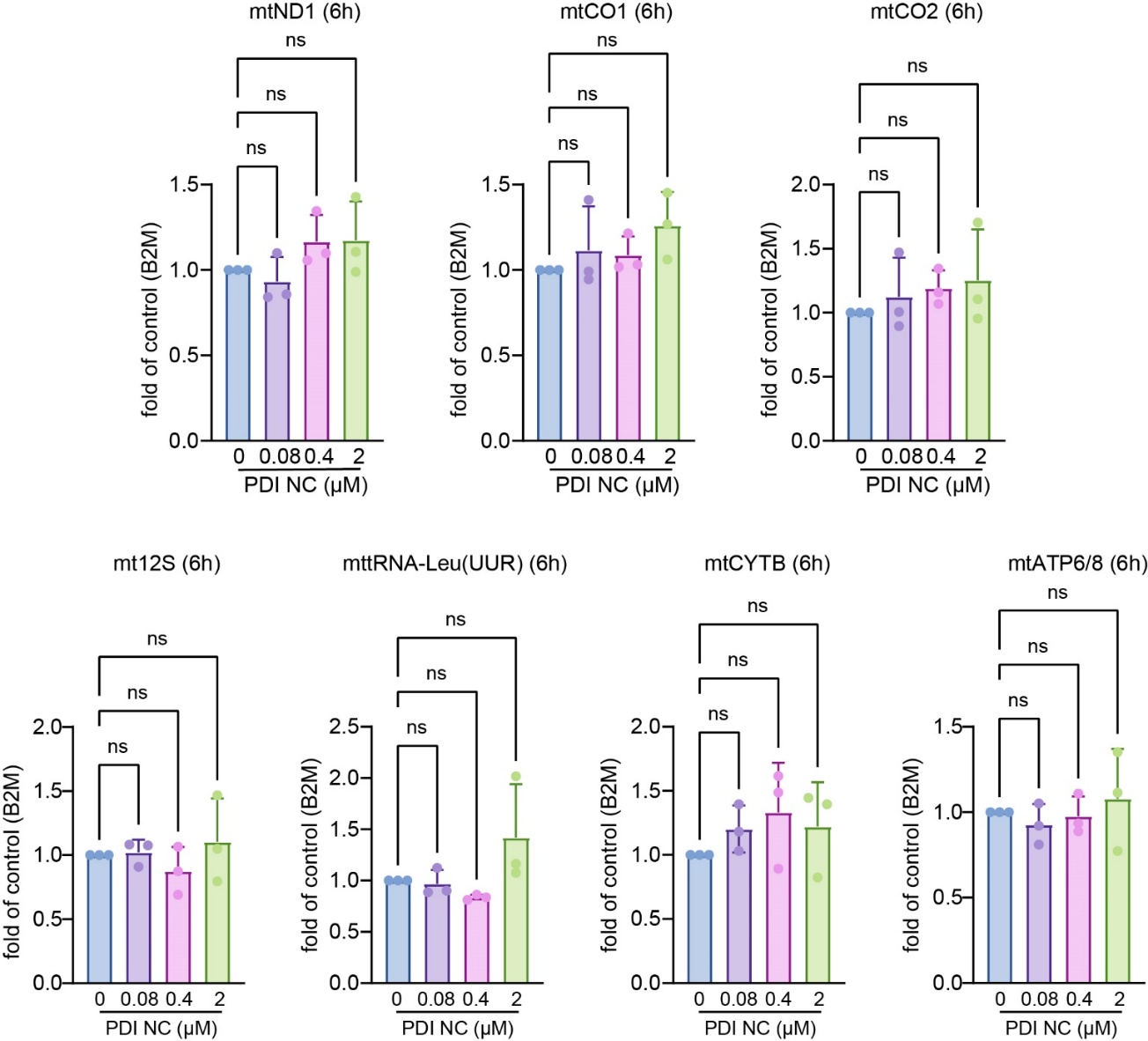

**Figure S27.** **PDI-NC exhibits negligible mtDNA transcription inhibition in MCF-7 cells.** MCF-7 cells were treated with PDI-NC at 0, 0.08, 0.4, or 2 µM for 6 h. Total RNA was extracted, and transcript levels of mtND1, mtCO1, mtCO2, mt12S, mttRNA-Leu (UUR), mtCYTB and mtATP6/8 were quantified by qRT-PCR using B2M as an internal control. Data are presented as fold change relative to untreated control (mean ± SD, n = 3). Statistical comparisons were made against the 0 µM group: ns, not significant. Overall, PDI-NC showed negligible transcriptional inhibition across most mitochondrial genes tested.

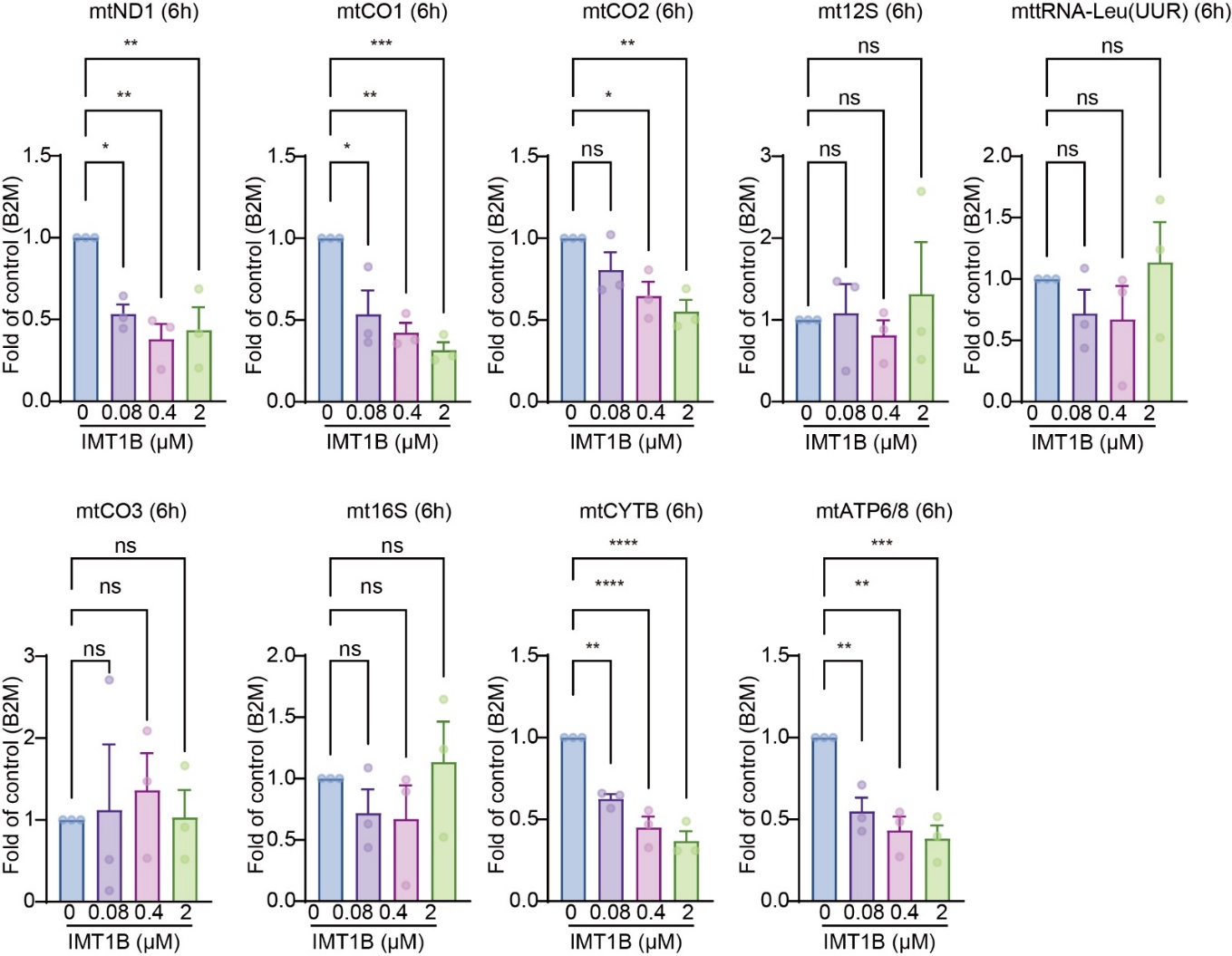

**Figure S28.** **IMT1B suppress mtDNA transcription in MCF-7 cells**. MCF-7 cells were treated with IMT1B at 0, 0.08, 0.4, or 2 µM for 6 h. Total RNA was extracted and transcript levels of mtCO1, mtCO2, mt12S, mt tRNA-Leu (UUR) mtCO3, mt16S, and mtCYTB, mtATP6/8 were quantified using B2M as the internal control. Data are presented as fold change relative to untreated control (mean ± SD, n = 3). Statistical comparisons were made against the 0 µM group: **p* < 0.05, ***p* < 0.01, ****p* < 0.001, *****p* < 0.0001; ns, not significant.

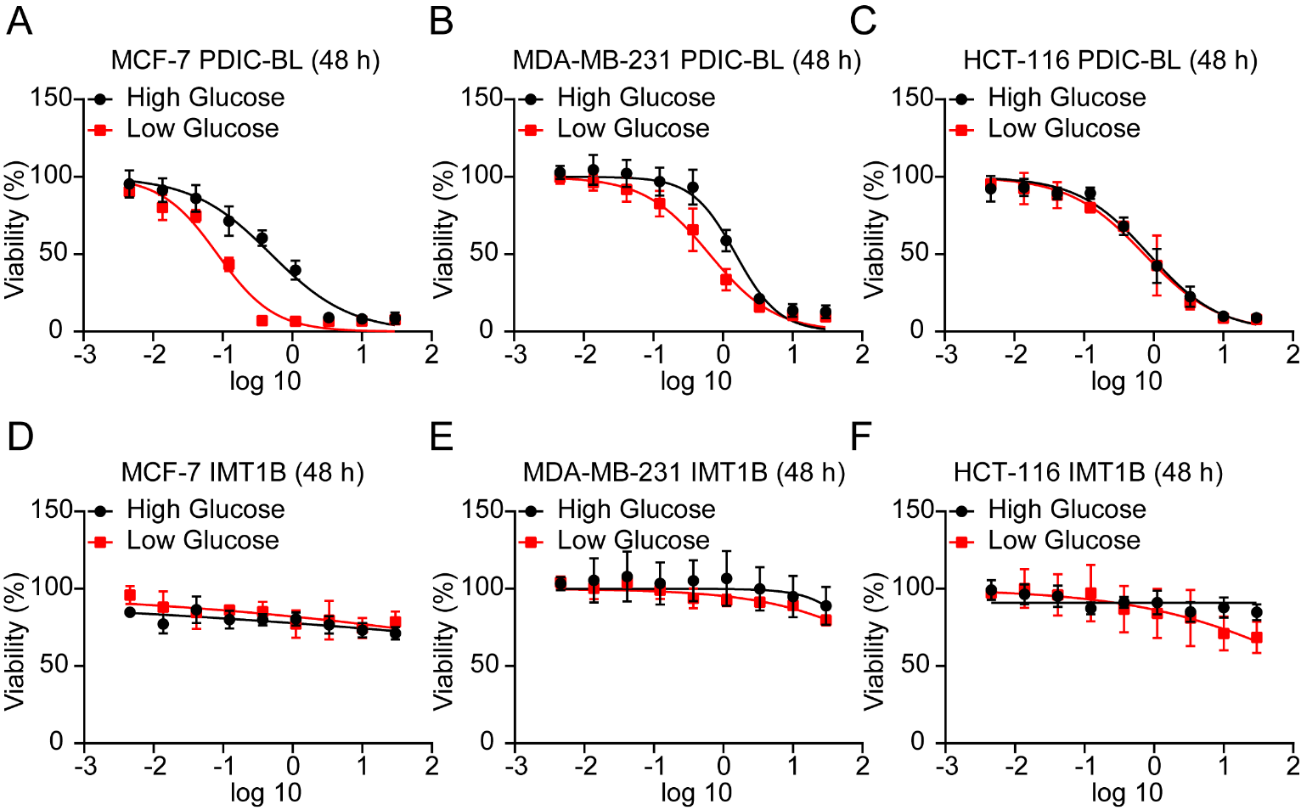

**Figure S29. Enhanced cytotoxicity of PDIC-BL under low-glucose condition in cancer cells.** (A-C) Dose-response curves for MCF-7 (A), MDA-MB-231 (B), and HCT-116 (C) cells treated with PDIC-BL for 48 h in either high-glucose (DMEM with 4500 mg/L glucose) or low-glucose (DMEM with 1000 mg/L glucose) medium. Under low-glucose conditions, the cytotoxicity of PDIC-BL was cell-line dependent. A significant enhancement in toxicity was observed in MCF-7 and MDA-MB-231 cells, with the IC_50_ values decreasing from 0.57 ± 0.06 µM to 0.11 ± 0.02 µM and from 1.15 ± 0.09 µM to 0.6 ± 0.2 µM, respectively. In contrast, the cytotoxicity in HCT-116 cells was not significantly altered, as the IC_50_ value shifted from 0.9 ± 0.4 µM in high-glucose medium to 1.0 ± 0.6 µM in low-glucose medium. (D-F) Dose-response curves for the same cell lines treated with IMT1B for 48 h under high- and low-glucose conditions, showing no significant change in potency. Even at the highest tested concentration of 30 µM IMT1B, MCF-7 and MDA-MB-231 cells maintained over 80% viability, suggesting that IMT1B exhibits weak antiproliferative activity in these cell lines (IC₅₀ > 30 µM). Data are presented as mean ± SD (n = 3).

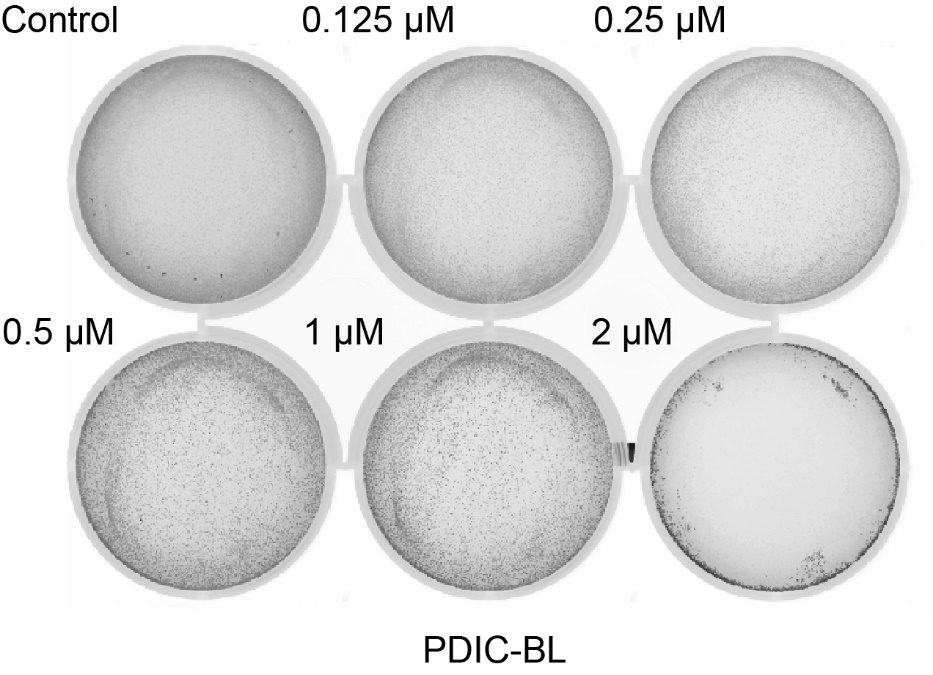

**Figure S30.PDIC-BL induces dose-dependent ROS generation in MCF-7 cells.** MCF-7 cells were incubated with PDIC-BL at 0 (Control), 0.125, 0.25, 0.5, 1, or 2 µM for 48 hours, followed by staining with DCFH-DA to detect intracellular ROS. Fluorescence image was obtained using Amersham Typhoon scanner. ROS-associated fluorescence intensity increased in a dose-dependent manner, reaching a maximum at 1 µM PDIC-BL. At 2 µM PDIC-BL, most cells were non-viable, resulting in a marked loss of signal due to compromised cell integrity.

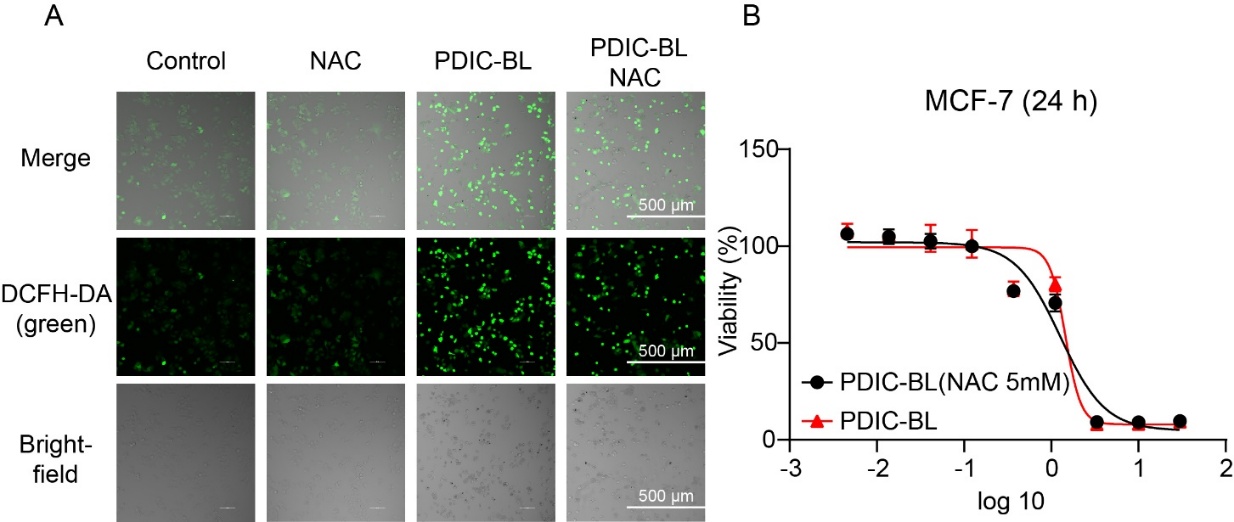

**Figure S31**. **NAC pretreatment does not attenuate PDIC-BL induced ROS production or cytotoxicity.** (A) Representative live-cell imaging of intracellular ROS in MCF-7 cells. Cells were pretreated with 5 mM N-acetylcysteine (NAC) or vehicle for 4 hours, followed by treatment with 10 µM PDIC-BL for 6 hours. ROS levels were detected using DCFH-DA fluorescence (green). NAC pretreatment failed to reduce ROS produced by PDIC-BL. (B) Dose-response curves for MCF-7 cell viability after PDIC-BL treatment for 24 hours, with or without a 4-hour pretreatment with 5 mM NAC. The calculated IC₅₀ were 1.2 ± 0.2 (with NAC) and 1.45 ± 0.02 µM (without NAC), indicating no significantly difference.

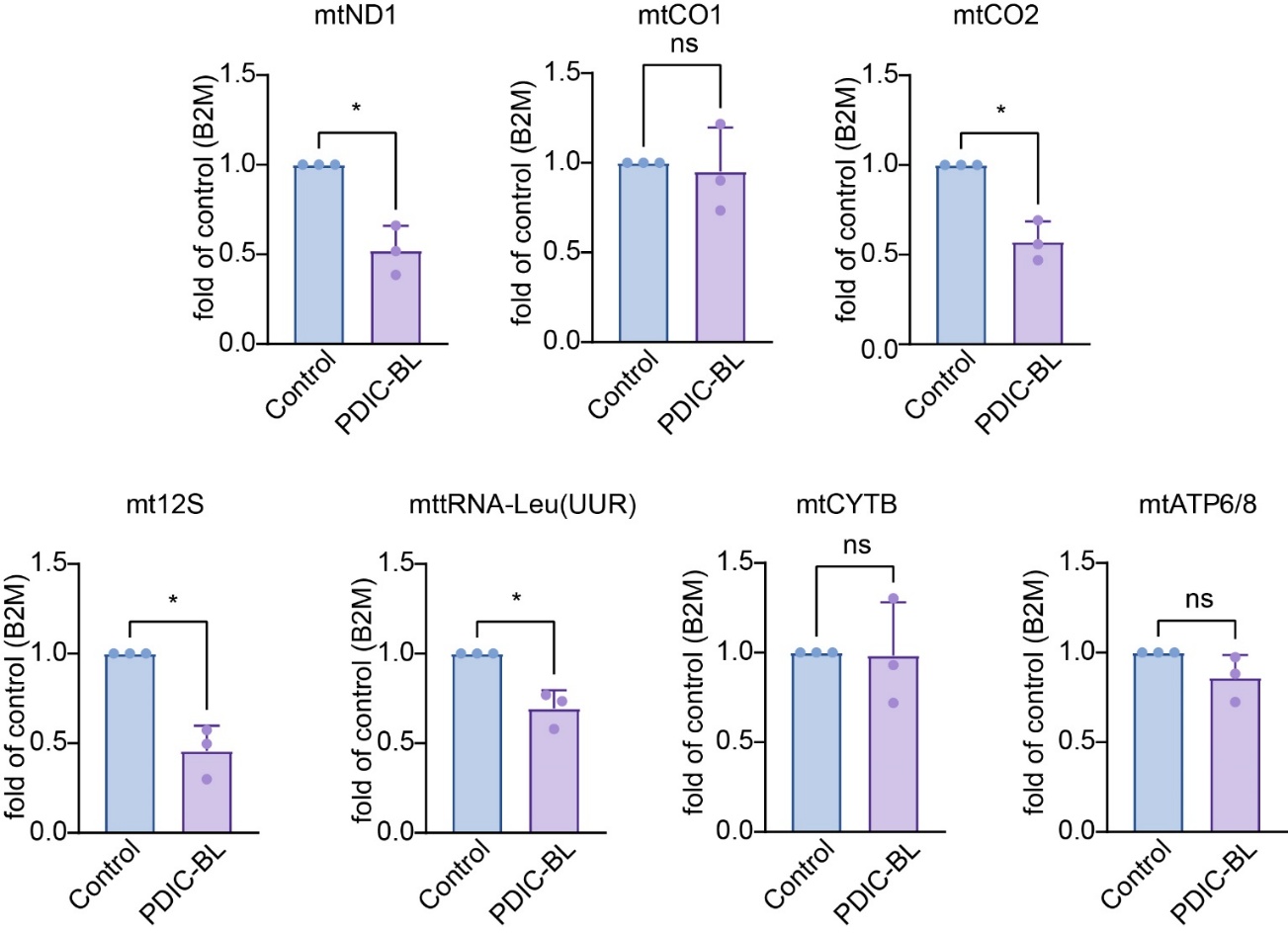

**Figure S32.PDIC-BL exhibits modest inhibition of mitochondrial gene transcription in vivo.** Quantitative RT-PCR analysis of mitochondrial gene expression in MCF-7 xenograft tumors from control and PDIC-BL–treated (2 mg/kg/day) nude mice. Total RNA was extracted, and transcript levels of mtND1, mtCO1, mtCO2, mt12S, mt-tRNA-Leu (UUR), mtCYTB, and mtATP6/8 were measured and normalized to B2M as an internal reference. PDIC-BL treatment resulted in a modest but detectable decrease in the expression of mtND1, mtCO2, and mt12S, while changes in other genes were not statistically significant. Data are presented as fold change relative to the control group (mean ± SD, n = 3). *p* < 0.05; ns, not significant.
